# Lineage-specific macrophage programs dictate metabolic suppression and stress responses associated with VBNC-like states in *Listeria monocytogenes*

**DOI:** 10.64898/2026.03.19.712867

**Authors:** Margherita Polidori, Geert Van Geest, Alba M. Neher, Camille Monney, Rémy Bruggmann, Anna Oevermann

## Abstract

*Listeria monocytogenes* (*Lm*) is a significant cause of central nervous system (CNS) infection in humans and animals, yet the mechanisms governing its intracellular lifestyle in brain-resident and infiltrating macrophages remain unclear. Using dual RNA sequencing combined with host proteomics, we map host–pathogen interactions in bovine microglia and monocyte-derived macrophages (MDMs), the two principal macrophage populations encountered by *Lm* in the CNS.

Although microglia and MDMs share core antimicrobial programs, they differ in their metabolic and immunological states, which dictate whether *Lm* adopts a replicative cytosolic lifestyle or a stress-tolerant intravacuolar state. In MDMs, nutrient restriction and phagolysosomal pressure drive metabolic suppression, robust stress and SOS responses, and induction of non-coding regulatory RNAs, shifting toward a dormant, viable but non-culturable phenotype. In microglia, the nutrient-rich cytosol supports bacterial growth, marked by upregulation of nucleotide salvage and carbohydrate and lipid metabolism. Functional analyses identify the stress-related genes *recA* and *rtcB* as contributors to intracellular persistence.

Together, our findings show that the fate of *Lm* is shaped not solely by canonical virulence genes but also by the interplay between bacterial stress adaptation and lineage-specific macrophage environments, highlighting macrophage ontogeny as a critical determinant of infection outcome.

**Authors Summary:** *Listeria monocytogenes* (*Lm*) is a deadly foodborne pathogen and major cause of central nervous system (CNS) infection (neurolisteriosis) in humans and ruminants, underscoring the close interdependence of animal health, food safety, and human health central to the One Health paradigm. During neurolisteriosis, *Lm* encounters distinct macrophage populations in the brain, yet how these cells shape bacterial behaviour has remained poorly understood. Here, we simultaneously captured the responses of *Lm* and bovine brain-resident microglia and infiltrating monocyte-derived macrophages (MDMs) during infection. We show that although both macrophage populations activate core antimicrobial programs, their metabolic and immune responses diverge profoundly, creating distinct intracellular environments that elicit different bacterial responses. In microglia, cytosolic bacterial replication is accompanied by transcriptional upregulation of growth-associated metabolic pathways. In contrast, MDMs impose nutrient limitation and phagolysosomal stress, triggering a global bacterial transcriptional reprogramming marked by metabolic shutdown, activation of stress and SOS DNA repair responses, and induction of regulatory non-coding RNAs. This response drives *Lm* into a dormant, stress-tolerant state that enables intracellular persistence despite immune pressure. Among activated genes, the bacterial recombinase A (RecA) and the RNA ligase RtcB are key contributors to bacterial persistence. Together, our findings reveal that macrophage ontogeny governs infection outcome by shaping both host and bacterial transcriptional programs, demonstrating that *Lm* persistence in the CNS depends not only on classical virulence factors but also on adaptive stress responses tuned to lineage-specific macrophage environments.

## Introduction

Despite the significant public health impact of bacterial infections in the central nervous system (CNS), the molecular mechanisms underlying these infections remain poorly understood ^1^. *Listeria monocytogenes* (*Lm*), a key model for the study of intracellular infection and host immunity ^2,3^, is a major cause of meningoencephalitis associated with high fatality rates ^4–7^. Its threat derives from *Lm*’s remarkable capacity to adapt to a wide range of intra-and extracellular conditions, transitioning from a saprophyte to a facultative intracellular pathogen capable of surviving in diverse environments, from food-processing environments to mammalian hosts.

This switch is driven by a finely tuned interplay between the stress activated sigma factor sigma B (SigB) and the master transcriptional regulator Positive regulatory factor A (PrfA), which orchestrates the expression of bacterial virulence factors required for cell entry, vacuolar escape, cytosolic replication, and cell-to-cell spread in phagocytic and non-phagocytic cells ^2^. SigB, under stress conditions such as gastrointestinal transit, initiates a stress response enhancing bacterial survival while priming PrfA activation ^8–10^. At body temperature, PrfA triggers the transcription of genes primarily located within the Listeria pathogenesis island 1 (LIPI-1) ^11,12^, encoding major virulence determinants including the internalins A and B (InlA, InlB) for host cell internalization, listeriolysin O (LLO) and phosholipases PlcA and PlcB for phagosomal escape, and the actin polymerizing protein ActA for autophagy avoidance and cell-to-cell spread ^13–17^. These factors collectively enable *Lm* to transition seamlessly from an environmental organism to an intracellular pathogen. Neurolisteriosis, a severe manifestation of foodborne *Lm* infection, causes intense CNS inflammation in humans and farmed ruminants^18,19^. In cattle, the disease manifests as rhombencephalitis, phenocopying brainstem infection in immunocompetent human patients ^18–20^. The inflammatory response in listerial rhombencephalitis is marked by microabcesses in cranial nerve nuclei and axonal tracts of the brainstem, populated by resident microglia, neutrophils and infiltrating monocyte-derived macrophages (MDMs) ^18,20,21^.

Emerging evidence suggests distinct roles for microglia and MDM in CNS diseases ^22,23^, including neurolisteriosis ^24,25^. These differences parallel previous observations that *Lm*‘s intracellular fate depends on the activation state and subtype of the infected macrophage ^26,27^. During hematogenous infection, circulating monocytes act as Trojan horses, transferring *Lm* to the endothelial cells of the blood-brain barrier and thereby facilitating bacterial dissemination into the brain ^25,28^. Brain-resident microglia act as transient amplification niches for *Lm*, supporting intracytosolic replication, and drive neutrophil recruitment into the infected brain ^29–33^. Conversely, MDMs confine *Lm* to the phagolysosomal compartment, where *Lm* can adopt a dormant, viable but non culturable (VBNC) state, akin to neutrophils and epithelial cells ^33–36^. This sophisticated dual intracellular survival strategy enables *Lm* to evade immune defenses and persist despite antibiotic therapy ^37,38^. The molecular mechanisms that dictate these cell-specific infection outcomes remain poorly defined.

Dual RNA sequencing (dual RNA-seq) is a powerful method that enables precise dissection of host-pathogen interactions by simultaneously capturing transcriptional responses from both organisms ^39,40^. Although this approach has illuminated the intracellular transcriptomes of several bacteria ^41–48^, its application to Gram-positive bacteria such as *Lm* has been limited due to technical challenges, including the low bacterial RNA yield relative to host RNA and the organism’s rigid cell wall hindering the extraction of bacterial RNA ^10,49,50^.

Here, we combine bulk dual RNA-seq with proteomics to study the molecular host-pathogen interactions that are involved in the cell-specific fate of *Lm* in two key CNS phagocyte populations: bovine primary microglia and MDMs. Our data uncovers distinct yet overlapping transcriptional landscapes pre- and post-infection that correspond to divergent infection phenotypes in macrophage lineages. In both host and bacterium, transcriptional responses lag phenotypic divergence, indicating that post-transcriptional regulation and intrinsic host cell factors shape bacterial fate, likely linked to the ontogeny of microglia and MDMs. Within the hostile intraphagosomal environment of MDMs, *Lm* downregulates core metabolic pathways, upregulates regulatory non-coding RNAs, and induces stress-related genes that coincide with the formation of a VBNC subpopulation. Among these, *recA* and the RNA splicing ligase *rtcB* emerge as candidate mediators of *Lm* persistence under macrophage-induced stress. Together, these findings provide an integrated view of *Lm*’s adaptive strategies within phagocytes, revealing cell-type specific mechanisms that shape infection outcomes and expose potential therapeutic vulnerabilities.

## Results

### Temporal profiling of bacterial and host responses in *Lm*-infected MDMs and microglia

To dissect the molecular pathways involved in the divergent intracellular fates of *Lm* in bovine MDMs and microglia, we performed simultaneous host-pathogen transcriptomic profiling combined with host proteomic analysis. Early during infection (45 minutes post-infection, pi), *Lm* exhibited a similar phenotype in both cell types (Fig 1A), as previously reported ^33^. By 6 hours pi, however, distinct infection outcomes emerged: in MDMs, *Lm* remained mainly confined to vacuoles, while in microglia, bacteria had escaped into the cytosol, as indicated by actin-polymerization (Fig 1A) ^33^. Vacuolar escape, which typically occurs around 30 minutes pi, is a key step in *Lm*’s intracellular lifecycle ^51^. Based on these dynamics, we selected 45 minutes and 6 hours pi as representative time points for dual RNA-seq analysis to compare microglia and MDM infection (Fig 1B-C). Since no substantial host response or major differences in bacterial gene expression between infected cell types were observed at 45 minutes pi, proteomic analysis was focused on the 6-hour time point. Mock-infected cells from the same animals served as controls for the host response, while bacteria grown for 6 hours in FBS served as a proxy for the extracellular infection stage (Fig 1B-C). This experimental design allowed us to capture the temporal dynamics of host and bacterial adaptations across distinct intracellular niches.

**Figure 1.**
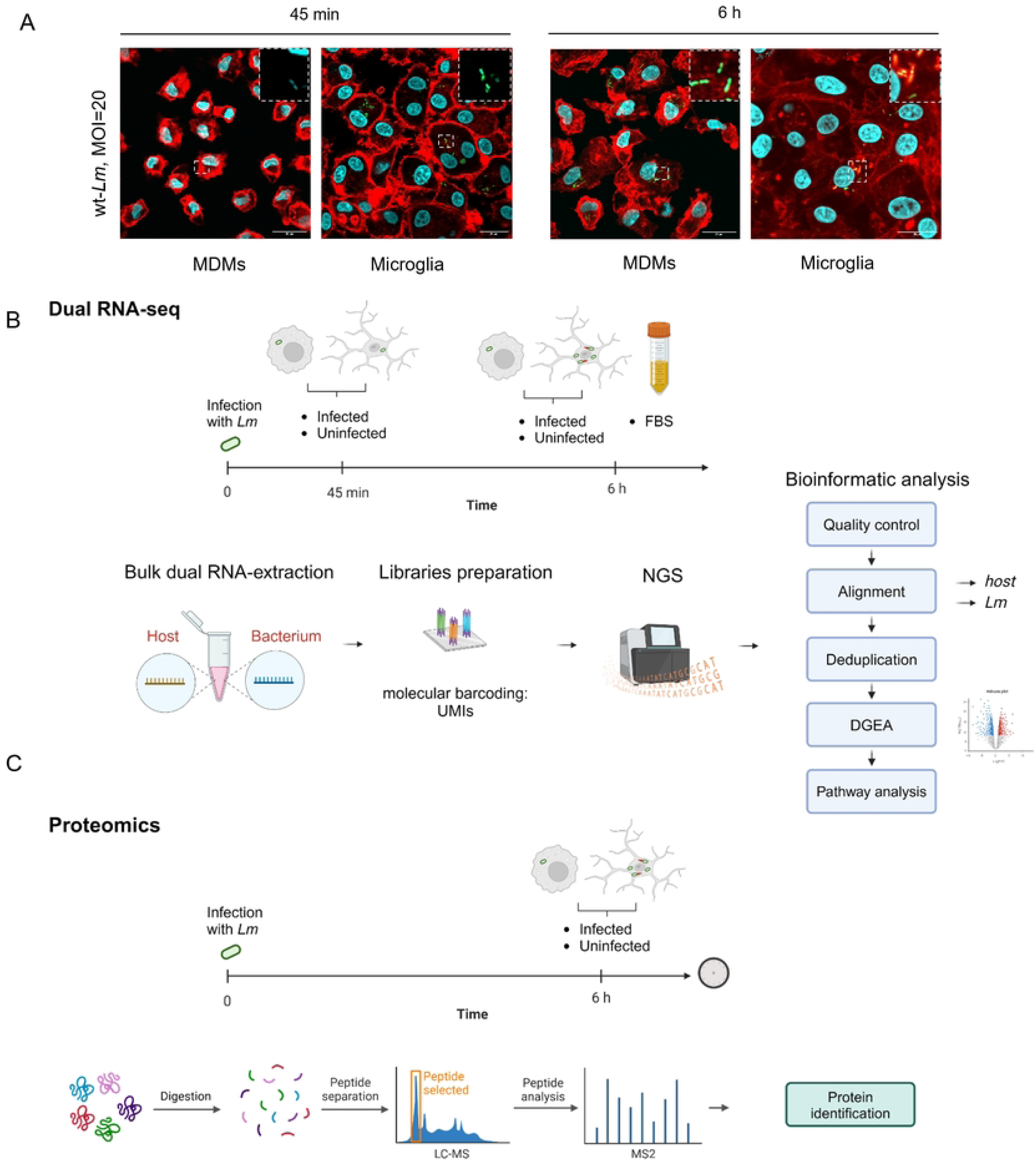
Experimental design for dual RNA-seq and proteomics analysis of host-pathogen responses in *Lm*-infected microglia and MDMs. (A) Immunofluorescence images illustrating phenotypic differences in *Lm* infection between MDMs and microglia in a gentamicin assay. *Lm* is visualized in green (wt-*Lm*-GFP), actin filaments in red (phalloidin), and nuclei in cyan (DAPI). While the phenotype of intracellular *Lm* appears similar at 45 min pi across macrophage subtypes, by 6 hours *Lm* polymerizes actin in microglia (indicating cytosolic localization; inset), whereas no actin polymerization is observed in MDMs (indicating intravacuolar localization). (B) Overview of the bulk dual RNA-Seq pipeline for infected bovine primary MDMs and microglia (upper panel). Cells were infected with a clinically relevant *Lm* CC1 strain (JF5203, wt-*Lm*) isolated from a bovine case of rhombencephalitis using a gentamicin protection assay at a MOI of 20. In parallel, extracellular wt-*Lm* was grown in FBS for 6 hours. Bulk dual RNA seq was performed on infected bovine primary microglia (n=3), MDMs (n=4), and extracellular wt-*Lm* (n=3), with mock-infected MDMs and microglia serving as controls for host transcriptomics. To enrich bacterial RNA, infected host cells were lysed in lysis buffer supplemented with lysozyme, and RNA was extracted at 45 minutes and 6 hours post infection. Following quality control, cDNA libraries were prepared using barcoded libraries constituted by Unique Molecular Identifiers (UMIs). Sequencing was performed on an Illumina platform, and bioinformatic analysis included quality control, deduplication, alignment to host and bacterial genomes, Differential Gene Expression Analysis (DGEA), and pathway analysis using Gene Set Enrichment Analysis (GSEA). (C) Simplified workflow for quantitative untargeted liquid chromatography-tandem mass spectrometry (LC-MS/MS) analysis of infected and mock-infected bovine primary microglia and MDMs (n=3 per condition). Cells were subjected to a gentamicin protection assay where wt-*Lm* were added at an MOI of 10. At 6 hours pi cells were harvested and digested for peptide separation and analysis. After data normalization, differential protein expression analysis was run across conditions, followed by pathway analysis. Illustration in A and B was created in BioRender (https://BioRender.com/ u24q725).

### *Lm* rapidly and dynamically adapts its transcriptome to disparate intracellular and extracellular environments

PCA of *Lm* transcriptomes revealed a clear separation between extracellular bacteria grown in FBS and those residing within MDMs or microglia (Fig 2A), indicating rapid transcriptional reprogramming upon phagocytosis. Intracellular *Lm* continued to adjust its gene expression between time points (Fig 2A-G). Notably, at 6 hours pi, the transcriptomic profiles of intracellular *Lm* segregated between MDMs and microglia, reflecting niche-specific adaptation (Fig 2A, C).

**Figure 2.**
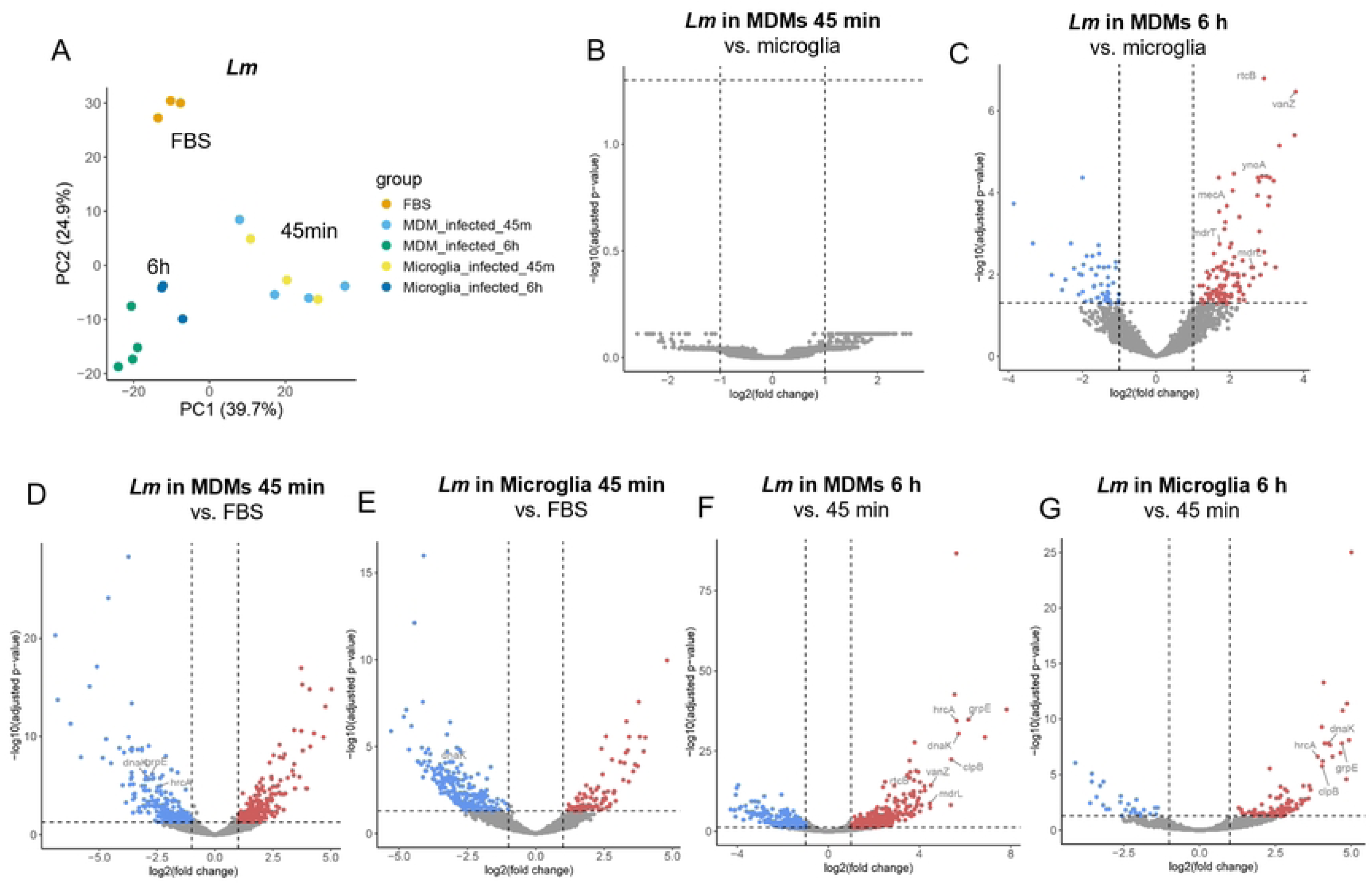
***Lm* transcriptome dynamics in bovine MDMs and microglia.** (A) PCA of the 500 most variably expressed bacterial genes in extracellular *Lm* grown in FBS for 6 hours and intracellular *Lm* in MDMs and microglia at 45 minutes and 6 hours pi. Different colors represent experimental groups, with each dot corresponding to a single biological replicate from an individual animal or bacterial culture. (B-G) Volcano plots illustrating differentially expressed *Lm* genes comparing intracellular conditions between phagocytes (B, C), intracellular to extracellular conditions (D, E) and across infection time points in MDMs (F) and microglia (G). Dotted lines indicate significance thresholds (|log2FC| ≥1; padj <0.05). Transcripts significantly upregulated in the first condition (bold) versus the second are highlighted in red, while those downregulated are shown in blue. (B) Differential expression of *Lm* transcripts in infected MDMs (right) and microglia (left) at 45 minutes. (C) Differential expression of *Lm* transcripts in infected MDMs (right) and microglia (left) at 6 hours. (D) Comparison of intracellular *Lm* transcriptome in early MDMs infection (right) to transcriptome of FBS grown *Lm* (left). (E) Comparison of intracellular *Lm* transcriptome in early microglia infection (right) to transcriptome of FBS grown *Lm* (left). (F) *Lm* transcriptome comparison between MDMs at 6 hours pi (right) and at 45 minutes pi (left). (G) *Lm* transcriptome comparison between microglia at 6 hours pi (right) and 45 minutes pi (left).

Differential gene expression (DGE) analysis revealed that *Lm* rapidly responds to the intracellular milieu, regulating 233 genes in MDMs and 136 in microglia within 45 minutes (Fig 2D-E, S2 Table). In both phagocyte types, bacterial transcriptional profiles continued to evolve over the 6-hour infection period. The response to infection involved a robust upregulation of stress genes, including the chaperones *clpE* and *clpB*, and heat shock proteins, such as *groL, groS, grpE, hrcA, dnaK*, and *dnaJ* (Fig 2F-G, S4A-B Fig), in line with previous studies in macrophages^52^. These findings indicate that *Lm* encounters stressful conditions in both phagocyte types, engaging in a core stress response shared across intracellular niches.

Additionally, at late stage of infection, *Lm* upregulated core virulence genes required for intracellular survival in both macrophage types, including members of the LIPI-1 (*hly, plcA, plcB, actA, mpl*) and additional virulence factors (*inlA, inlB, inlC, uhpT*) (Fig 2, Fig 3A, Suppl.Table 3). In contrast, the major transcriptional regulators PrfA and SigB, the SigB-dependent target OpuCA ^9^, and LLS A and B showed no marked transcriptional dynamics, although some exhibited a non-significant trend toward higher expression in FBS-grown bacteria compared with intracellular stages. Similarly, the two internalin genes *inlA* and *inlB* were expressed at higher levels in FBS-grown bacteria than intracellularly at 45 minutes of infection (Fig 3A, S3 Table), unlike *plcA, plcB, actA, mpl*, *inlC* and *uhpT,* which were expressed at low levels in FBS and robustly upregulated at 6 hours pi in both microglia and MDM. These distinct expression patterns likely reflect *sigB* regulation of *inlA* and *inlB* ^53^ and the influence of glutathione availability in both phagocytes ^8,54–56^. Notably, the expression of the major virulence genes (e.g. *prfA, hly, plcB, plcA, actA, mpl, inlA, inlB, sigB*) was comparable between microglia and MDMs at both time points (Fig 2, Fig 3A, S3 Table), indicating that differences in canonical virulence gene expression do not account for the distinct intracellular phenotypes observed (Suppl. Table 3).

**Figure 3.**
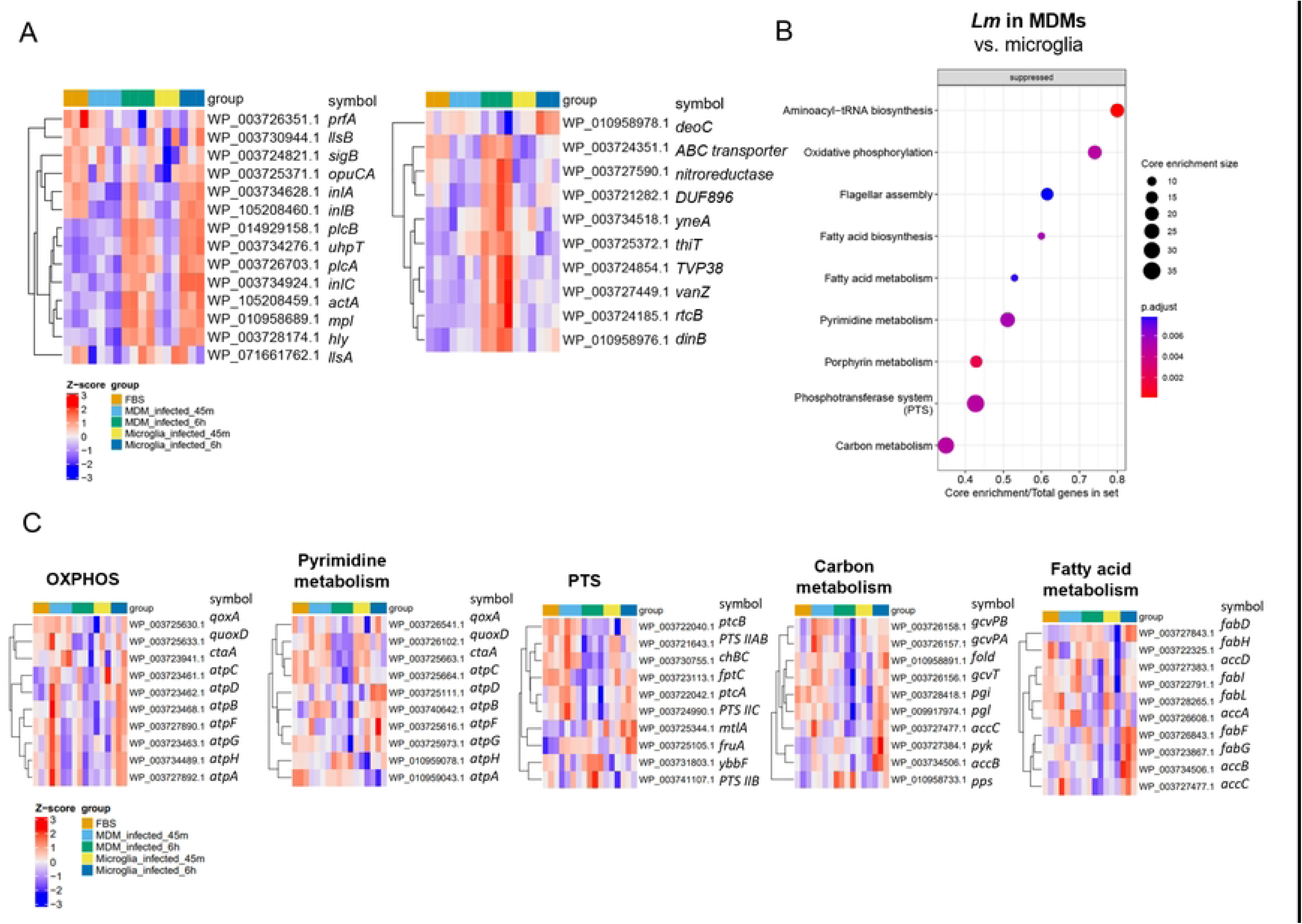
Adaptatation of *Lm* gene expression to different intraphagocytic environments. (A-B) Heatmaps illustrating *Lm* transcriptional adaptation to FBS and the intracellular environment in MDMs and microglia across time points. A: Heatmap of *Lm* core virulence genes previously characterized as essential for its intracellular survival. B: Heatmap of the top 10 significant DEGs (|log2FC| ≥1; padj<0.05) in *Lm* infecting MDMs compared to microglia. The Z-score represents scaled log-normalized counts. The gene symbols refer to the annotation of the NCBI Reference Sequence: NZ_LT985474.1. (C) Dot plot of the top 10 KEGG pathways significantly suppressed in *Lm* (FDR < 0.05) infecting MDMs compared to microglia at 6 hours pi (qvalue<0.05). (D) Heatmaps displaying the top 10 *Lm* transcripts (gene symbols) associated with key metabolic pathways: Oxidative phosphorylation (OXPHOS, map00190), pyrimidine metabolism (map00240), phosphotransferase system (PTS, map02060), carbon metabolism (map01200), and fatty acid biosynthesis (map00061).

Despite these shared responses, transcriptional divergences between intracellular *Lm* in MDMs and microglia became increasingly evident over time, with 170 genes differentially expressed at 6h pi: 121 upregulated in MDMs and 49 in microglia (Fig 2B-C, S2 Table). Genes robustly upregulated in MDMs included those associated with stress, including *vanZ, rtcB, yneA, DUF896, dinB,* and metabolic adaptation, including ABC transporter, a nitroreductase family protein, and the thiamine transporter *thiT* (Fig 3B). This transcriptional pattern suggests that MDMs impose a strongly hostile and stress-inducing environment, consistent with the observed prolonged vacuolar confinement ^33^. In contrast, cytosolic *Lm* in microglia mounted a far less pronounced stress response and strongly upregulated the deoxyribose-phosphate aldolase *deoC* (Fig 3B), a key enzyme in nucleoside and pyrimidine salvage, indicating a shift toward metabolic adaptation and nucleotide acquisition to support rapid replication.

In summary, *Lm* exhibits exceptional transcriptional plasticity, switching to a stress-resistant and survival-oriented persister state in the hostile environment of the MDM’s vacuole, while shifting to a metabolically active, replicative program in the permissive cytosol of microglia. This dual strategy maximizes bacterial survival across distinct intracellular niches.

### *Lm* fine-tunes metabolism for intracellular adaptation in MDMs and microglia

Intracellular *Lm* rely on diverse metabolites to support catabolic and anabolic processes, including carbon sources such as PTS-transported glucose, mannose and chitobiose, as well as glycerol, hexose phosphates and L-glutamine ^54,57–59^. During infection, *Lm* encounters fluctuating nutrient availability across host cell types and subcellular compartments. Accordingly, we observed both shared and niche-specific metabolic adaptations in *Lm* infecting microglia versus MDMs. Conserved metabolic responses in both macrophage subtypes included upregulation of the G1P transporter gene *uhpT* (Fig 3A, S3 Table), which facilitates host-derived hexose phosphate uptake and is essential for intracellular infection ^60,61^. In contrast, several bacterial metabolic pathways were differentially regulated in the two cell types. In MDMs, *Lm* suppressed, whereas in microglia it activated, key metabolic pathways, including aminoacyl-tRNA biosynthesis (map00970, indicating reduced translation and growth), oxidative phosphorylation (OXPHOS, map00190) and pyrimidine metabolism (map00240) (Fig 3C, S3 Table). At 6-hour pi, *Lm* in MDM decreased the expression of genes associated with the phosphotransferase system (PTS, map02060), indicating reduced carbohydrate uptake, fatty acid biosynthesis (map00061) and metabolism (map01212), carbon metabolism (map01200), while upregulating valine, leucine and isoleucine biosynthesis (map00290) as compared to microglial cells (Fig 3C, S3 Table). This transcriptional signature is consistent with a shift in MDMs towards an energy-conserving, non- to low-replicative, persister-like state.

Within OXPHOS, *Lm* downregulated *qoxA*, *qoxD* and *ctaA*n in MDMs (Fig 3C) over the course of the infection (Fig 3D, S3 Table). Notably, *qoxD*, encoding a subunit of the quinol oxidase complex ^62^, was significantly lower at 6h compared to *Lm* in microglia and to the 45-minute time point (Fig 3D, S3 Table).

In pyrimidine metabolism, *Lm* downregulated de-novo synthesis genes *pyrE*, *pyrF* and *pyrG* ^63,64^ in both macrophage types at 6 hours pi (Suppl. Table 3). However, *pyrF* expression remained higher in microglial cells at 6 hours pi than in MDM (Fig 3D, S3 Table), suggesting enhanced host-derived nucleotide salvage in the nutrient-rich cytosol compared to *Lm* in MDM, where reduced expression may reflect slowed replication.

Also, *dnaX*, encoding γ and τ subunits of the DNA polymerase III for restart of stalled replication forks ^65,66^, decreased in both macrophage types upon entry but then rebounded by 6 hours pi (Fig 3D, S3 Table), indicating a transient replication slowdown followed by restart. In contrast, *dnaN,* essential for DNA replication, trended towards downregulation in MDM and upregulation in microglia at 6 hours pi, causing a relative enrichment in microglia at 6h (Fig 3D, S3 Table). This pattern suggests that replication restart mechanisms (*dnaX*) are retained in both environments, but efficient DNA synthesis (*dnaN*) is favored in the microglial cytosol and restricted within MDM phagosomes. In microglia, *Lm* also maintained *pdp* upregulation, encoding a pyruvate dehydrogenase phosphatase activating the pyruvate dehydrogenase complex, and strongly induced *fruK*, encoding a 1-phosphofructokinase, consistent with enhanced oxidative metabolism and fructose catabolism in the cytosol, feeding the tricarboxylic acid (TCA) cycle (Fig 3D, S3 Table). In contrast, *pdp* expression in MDMs returned to extracellular levels after early upregulation and *fruK* was not induced (Fig 3C, S3 Table), indicating a shift away from PDH- and TCA-linked metabolism under nutrient restriction.

In the PTS pathway, genes including *fruA*, *mtlA*, and *chbC*, and *ybbF* exhibited divergent expression 6h pi (Fig 3D, S3 Table). *Lm* upregulated the fructose transporter encoding gene *fruA* ^62^ in both macrophage types, but more strongly in microglia. The mannitol transporter *mtlA* was clearly upregulated only in microglia, while the phosphoenolpyruvate-protein phosphotransferase *ptsI* increased in microglia during infection without reaching significance relative to MDMs (Fig 3D, S3 Table). In contrast, the chitobiose transporter encoding gene *chbC* ^62^ was selectively downregulated in MDM, along with other genes including *ptcA*, *ptcB*, *ulaA* and *fptC* (Fig 3 C, S3 Table). Conversely, *ybbF, encoding* the PTS transporter subunit EIIC was the only PTS gene strongly induced in MDM but not in microglia. Although early studies suggested that the downregulation of PTS systems in macrophages makes them dispensable for intracellular survival ^67–69^, recent evidence supports their critical role in intracellular growth and virulence ^54^, in line with our observations.

Carbon metabolism genes of *Lm* that exhibited niche-specific dynamics in infected microglia and MDM included *gcvpA, gcvpB* and *gcvT* (Fig 3D, S3 Table), components of the glycine cleavage system ^70^, which showed early induction in MDMs followed by repression while remaining relatively stable in microglia (Fig 3D, S3 Table). Late-stage MDM infection was further characterized by downregulation of *pgi* and *pgl,* involved in glycolysis and the pentose phosphate pathway. In microglia, *pgi* and *pgl* expression dropped early but recovered at 6 hours pi (Fig 3D, S3 Table). Although not differentially expressed between niches, *pyk* was upregulated in microglia at 6h, whereas *pps*, encoding phosphoenolpyruvate synthase for gluconeogenesis, increased in MDM, indicating activation of gluconeogenesis (Fig 3D, S3 Table). Together, these transcriptional signatures suggest that *Lm* sustains glycolytic and TCA-linked metabolism to support replication in the microglial cytosol, while in nutrient-restricted MDM phagosomes *Lm* suppresses glycolysis and engages gluconeogenesis and transient glycine catabolism. This is consistent with a metabolic shift from carbohydrate catabolism to carbon recycling under phagosomal nutrient stress ^71^.

Fatty acid biosynthesis pathways were also differentially regulated between *Lm* infecting MDMs and microglia. Acetyl-CoA carboxylase subunit genes (e.g. *accB* and *accC*) and type II fatty acid synthesis (FASII) genes (*fabD, fabG* and *fabF)* ^72^ were upregulated in microglia (Fig 3D, S3 Table), while *fabI* and *fabL* were downregulated in MDMs (Fig 3D, S3 Table), suggesting increased demand for membrane synthesis during active replication in microglia ^73^.

Overall, these data reveal that *Lm* adapts its metabolism in a niche-specific fashion. The nutrient-rich microglial cytosol supports active replication by engaging host nucleotide salvage, oxidative and carbohydrate metabolism, and DNA replication. In contrast, within the nutrient restricted phagosomes of MDM, *Lm* supresses biosynthetic and energy-producing pathways in favor of a stress-tolerant, low-replication state, reflecting divergent survival strategies in permissive versus restrictive intracellular environments.

### Niche-specific regulation of *Lm* noncoding small RNAs during infection

Niche-adaptive metabolic and stress responses were also reflected in *Lm*‘s small (s)RNA signature. While *Lm* clearly adapted to the intracellular environment in both macrophage types, the early bacterial sRNA response largely overlapped between microglia and MDMs S3 Fig A-B), mirroring the similarity observed in the coding transcriptome at this stage (Fig 2A-B). Significant divergence emerged only at 6h pi, when the sRNA response was stronger in MDMs.

The *Lm* riboswitch *rli55, which* modulates *eut* genes involved in ethanolamine metabolism and is relevant for *Lm* virulence ^74^, was strongly upregulated at 45 minutes in both macrophage types (S3 Fig C, S6 Table) and repressed at 6h (S3 Fig D, S6 Table). This pattern suggests a conserved early stress-adaptation program, with ethanolamine likely serving as an alternative carbon and nitrogen source during initial intracellular adaptation.

T-box riboswitches, including T-boxes 4, 5, 7, were strongly upregulated upon entry in both macrophage types and then further increased in MDMs, while they stabilized in microglia. *Lm* further upregulated T-box 2 and 8 in MDM at 6h, while it downregulated T-boxes 3 and 8 in microglia at this later stage (S6 Table). This led to relative enrichment of multiple T-boxes (3, 4, 6, 7, 8) in MDMs (S3 Fig B, S6 Table), suggesting their strong induction as part of a metabolic adaptation to amino acid limitation and the accumulation of uncharged tRNAs in response to suppression of the aminoacyl-tRNA biosynthesis pathway in the intravacuolar environment of MDMs (Fig 3B) ^75,7663,64^. In microglia, the transient early T-box pulse likely reflects initial nutrient limitation prior to cytosolic escape. Consistent with our findings, T-box riboswitch expression has previously been reported during *Lm* infection of murine macrophages ^10,75,79^. The late-stage upregulation of *rli53*, an antibiotic- and pressure-responsive riboregulator, in MDM but not in microglia, further supports the idea of a sustained niche-specific stress response and suggests a broader role for this riboswitch in stress tolerance ^80,81^. Likewise, *rli24*, an uncharacterized intergenic sRNA of unknown function, was strongly and specifically induced in MDM, suggesting a role in stress adaptation.

Collectively, these results highlight *Lm*’s ability to dynamically regulate its metabolism and sRNA expression in response to the distinct metabolic and stress conditions encountered within host cell niches, optimizing survival and replication across various host environments.

### *Lm* activates a strong stress response to ensure intracellular survival in MDMs

Our analysis revealed that several bacterial genes related to stress and SOS response were strongly upregulated in MDMs during late infection, showing relative enrichment compared with microglia (Fig 4A, S3 Table). The most strongly induced gene encoded a VanZ family protein (Fig 4A, S3 Table). Genes enriched in the SOS response (GO:0006950, False Discovery Rate/FDR= 0.0223) included the transcriptional repressor *lexA*, the cell division suppressor *yneA*, the DNA recombinase *recA*, and *dinB,* encoding DNA polymerase IV. Stress response genes (GO:0009432, FDR= 0.0415) comprised *clpE, clpB, dnaJ, groL* (Fig 4A, Suppl. Table 3, S6 Fig), all known to be necessary for intracellular survival in both phagocytic and non-phagocytic cells ^58,82^. The DNA repair gene *uvrB,* which contributes to both stress and SOS response pathways, was also upregulated (Fig 4A, S3 Table, S6 Fig). Additional genes strongly upregulated in MDMs included *mecA*, encoding for an adaptor protein, *mdrT*, encoding for a cholic acid efflux major facilitator superfamily (MFS) transporter, *mdrL*, encoding for *a* multidrug efflux MFS transporter, and *rtcB*, encoding for an RNA-splicing ligase (Fig 4A, S3 Table, S6 Fig). Some stress-related genes (*dnaJ, clpB, clpE, groL, uvrB, and lexA*) were also upregulated in microglia over time and relative to FBS-grown controls (Fig 4A, S3 Table, S6 Fig). Moreover, chaperone and heat shock protein genes (*dnaK, grpE, hrcA,* and *clpB)* showed similar induction in both MDMs and microglia (Fig 4A, S3 Table, S6 Fig), indicating that both host cell types impose stress on *Lm*, though the response is more pronounced in MDMs. Overall, the enhanced activation of stress-related and SOS pathways in MDM-associated *Lm* aligns with the formation of VBNC bacteria reported in these cells and other bacterial species33,83.

**Figure 4.**
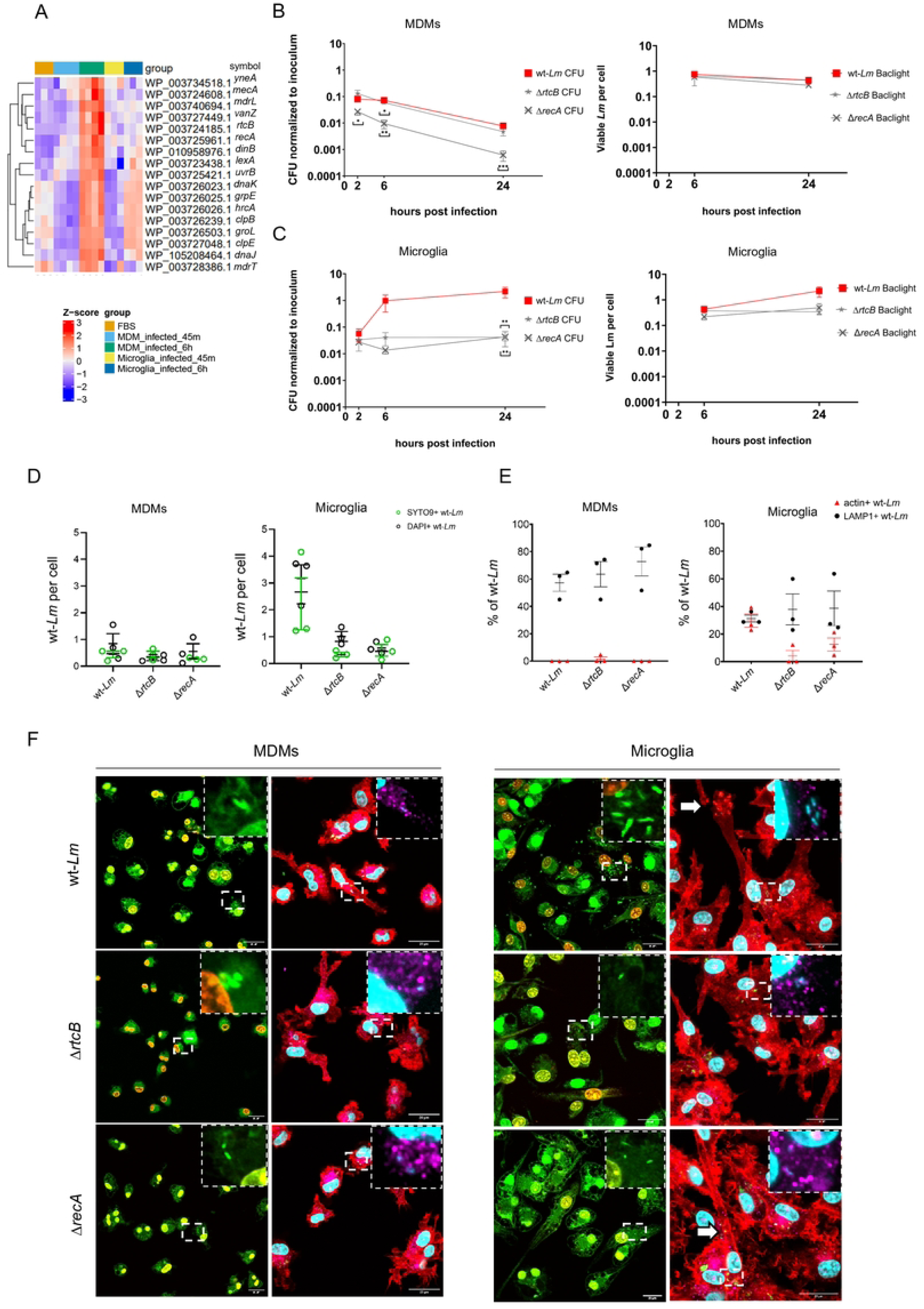
*Lm* mounts a stress response to the intracellular environment within MDMs. (A) Heatmap of selected *Lm* stress and SOS-response-related transcripts that are differentially expressed between phagocytes (|log2FC| ≥1; padj<0.05). The Z-score represents scaled, log-normalized counts. Gene symbols refer to the annotation of the NCBI Reference Sequence: NZ_LT985474.1. (B-F) Among the *Lm* stress and SOS-response genes identified in (A) and tested functionally, *recA* and *rtcB* affected the infection phenotype in gentamicin protection assays using bovine MDM and microglia. Results for the other tested genes, which did not show a significant effect on infection phenotype, are shown in Suppl. Fig. 6. Quantification of CFUs (B–C) were performed in three (Δ*recA*) or four (wt-*Lm*, Δ*rtcB*) independent experiments (three in technical triplicates (MDMs) and duplicates (microglia). Data are shown as mean ± SEM. Quantifications of bacteria in BacLight assays and IF (B–E) were obtained from three biological replicates each. (B) Intracellular CFUs normalized to the inoculum (left panel) and SYTO9+ viable bacteria per cell (BacLight assay, right panel) in MDMs infected with wt-*Lm* (control), Δ*rtcB*, and Δ*recA*. (C) Intracellular CFUs normalized to inoculum (left panel) and SYTO9+ bacteria per cell (BacLight assay, right panel) in microglia infected with wt-*Lm* (control), Δ*rtcB*, and Δ*recA*. Mann-Whitney test: * p<0.05, **p<0.01, ***p<0.0001. (D) Comparison between SYTO9+ viable bacteria (n=3) and DAPI stained number of total bacteria (n=3) per cell after 24 hours of infection. (E) Percentage of intravacuolar bacteria associated with LAMP1 or with polymerized actin (clouds or tails, indicating cytosolic location) in microglia and MDMs. Error bars indicate the standard error of the mean (SEM). (F) Confocal images of MDMs and microglia infected for 24 hours with wt-*Lm* (control), Δ*rtcB*, and Δ*recA*. In the left panel, live bacteria are labeled green with SYTO9, while dead or permeabilized bacteria are labeled in red with PI. The inset provides a higher-magnification view of the highlighted region. On the right panel, cells are stained with Phalloidin, LAMP1, DAPI and *Lm* antiserum. Actin tails and clouds are indicated by the white arrows. The inset provides a higher-magnification view of the highlighted region stained with DAPI and LAMP1.

To further explore the role of stress-response genes in intra-macrophage survival, we generated deletion mutants for selected upregulated stress-related genes identified (*mecA, vanZ, mdrT, yneA*, *rtcB*, and *recA,* Fig 4 A). Whole genome sequence analysis confirmed the intended deletions and identified a few off-target SNPs (S5 Fig, S7 Table). We assessed the infection phenotypes of deletion mutants in MDMs and microglia. Intracellular survival was quantified by CFU counts and microscopy, including quantification of SYTO9+ live bacteria in BacLight assays and DAPI + bacteria in immunofluorescence experiments (Fig 4B-D, S6A-G Fig).

All mutants displayed growth kinetics comparable to wt-*Lm* in nutrient-rich broth, indicating that the targeted genes are not required for basal growth or metabolism (S6A Fig). However, distinct intracellular phenotypes emerged upon infection of MDMs and microglia. In MDMs, Δ*mecA,* Δ*vanZ,* Δ*mdrT,* Δ*rtcB* behaved similarly to wt-*Lm,* while Δ*yneA* showed a moderate yet significant reduction in intracellular CFUs at 6 hours pi, and Δ*recA* showed up to a tenfold reduction across all time points, which was significant at 6 and 24h (Fig 4B, S6C Fig). Despite this, microscopy revealed that all mutants, including Δ*recA*, maintained SYTO9+ viable counts comparable to wt-*Lm* (Fig 4B, S6C Fig), confirming the presence of a substantial VBNC fraction in MDMs ^33^, which was further increased in Δ*recA*.

In microglia, the Δ*vanZ* mutant mirrored wt-*Lm* behaviour, while Δ*mecA,* Δ*mdrT* and Δ*yneA* showed reduced CFU levels that did not reach statistical significance at 24h (S6D Fig). By contrast, Δ*recA* and Δ*rtcB* displayed a marked reduction in recoverable CFUs from as early as 2h, culminating in a nearly two-log reduction at 24h (Fig 4C). Consistent with previous reports^33^, SYTO9+ viable counts of wt-*Lm* increased over time and closely paralleled CFU numbers in microglia (Fig 4 C). All mutants exhibited slightly reduced total bacterial numbers and SYTO9+ counts at 24h, though differences were not statistically significant (Fig 4B-D, S6D Fig). Quantification of bacteria per cell based on SYTO9+ staining aligned with the counts of DAPI positive bacteria obtained by IF microscopy (Fig 4D-F). These analyses revealed comparable intracellular bacterial loads for wt-*Lm*, Δ*recA,* and Δ*rtcB* in MDMs, but significantly reduced numbers for Δ*recA,* and Δ*rtcB* in microglia (Fig 4D). Notably, in microglia, Δ*rec*A and Δ*rtc*B exhibited high SYTO9+ bacterial numbers despite markedly reduced CFU recovery, mimicking the VBNC-like phenotype observed for wt-*Lm* in MDM. A similar reduction in intracellular CFUs for Δ*recA* and Δ*rtcB* was observed in the human microglial cell line HMC-3 at 24h pi (S6B Fig). This effect was less pronounced at early time points, with slightly higher intracellular colonies compared to wt-*Lm* at 2h pi, followed by a modest reduction of intracellular bacteria at 6h pi (S6B Fig).

As VBNC in MDMs localize predominantly within phagosomes, we compared the intracellular localization of Δ*rtcB*, and Δ*recA* mutants with wt-*Lm* in both cell types using actin polymerization (for intracytosolic localization) and LAMP1-association (for intravacuolar localization) (Fig 4E-F). As expected, actin-polymerizing bacteria were absent in MDMs. The LAMP1+ association was modestly higher in mutants (57% in wt-*Lm*, 64% in Δ*recA*, and 73% in Δ*rtcB)*, suggesting increased restriction to the phagolysosomal system (Fig 4E). In microglia, Δ*recA* and Δ*rtcB* displayed a trend to reduced cytosolic access (fewer actin tails) and increased LAMP1 association (Fig 4E), consistent with impaired vacuolar escape.

Taken together, these findings show that activation of stress adaptation pathways is a central strategy for bacterial survival within diverse intra-phagocytic environments. Although SOS and stress response genes are strongly induced in MDMs, they appear largely dispensable for intracellular viability under our conditions. By contrast, *rec*A or *rtc*B contribute to stress tolerance and VBNC induction. *Lm* does not build up a significant VBNC population in microglia unless *rec*A or *rtc*B are deleted, two genes that are highly expressed in MDM-associated bacteria during late infection. The reduced cytosolic presence of Δ*rtcB* and Δ*recA* mutants in microglia likely reflects impaired fitness in phagosomes limiting vacuolar escape and highlighting these genes as key determinants of the intravacuolar bacterial lifestyle.

Notably, deletion of *recA*, but not *rtcB*, increased the VBNC bacteria fraction in MDMs, suggesting the loss of *rtcB* may be compensated by broad upregulation of other stress-response genes identified in our transcriptomic data. In contrast, *recA*, a central player in the SOS response and DNA repair, appears non-redundant in MDMs, and its loss exacerbates VBNC formation. In microglia, where the bacterial stress response (including *recA* and *rtcB*) is less pronounced, deletion of either gene may tip the balance toward VBNC induction. Collectively, these findings suggest that *recA* and *rtcB* support intracellular persistence through niche-specific stress adaptation and that the activated stress-response network in MDM may provide resilience against single-gene deletions.

### Host transcriptional responses to *Lm* infection only partially overlap in MDMs and microglia

To validate the identity of our cultured macrophage populations, we assessed the expression of lineage-defining transcripts. Microglia expressed high mRNA levels of *P2RY12*, *SALL1* and *APOE* relative to MDMs, while MDMs showed elevated *F13A1*, *CD14*, and *CD63* (S8 Fig I), compatible with reported bovine MDM and microglia signatures ^84^. Notably, mock-infected microglia showed no transcriptional changes during culture, and MDMs exhibited only minimal alterations, ruling out transcriptomic artifacts due to experimental procedures (S8E-F).

PCA of host transcriptomes revealed clear segregation between MDMs and microglia, independent of infection status (Fig 5A). Among all conditions, only infected MDMs at 6 hours pi clustered distinctly from uninfected controls (Fig 5A). DGEA further confirmed robust transcriptomic differences between MDMs and microglia before and after infection, reflecting their divergent ontogeny (Fig 5C-E; S7 A-B; Fig 6) ^84^. Over the course of infection, MDM upregulated 781 genes and microglia 106, of which 71 were shared (S7 C-D Fig, S2 Table). At 6h, 1780 genes were upregulated in infected MDMs and 1530 in infected microglia (Fig 5E, S2 Table). In the uninfected state, 1271 and 1062 genes were differentially expressed in MDMs and microglia, respectively (Fig 5D, S2 Table), mirroring the late divergence observed in bacterial transcriptomes (Fig 2B).

**Figure 5.**
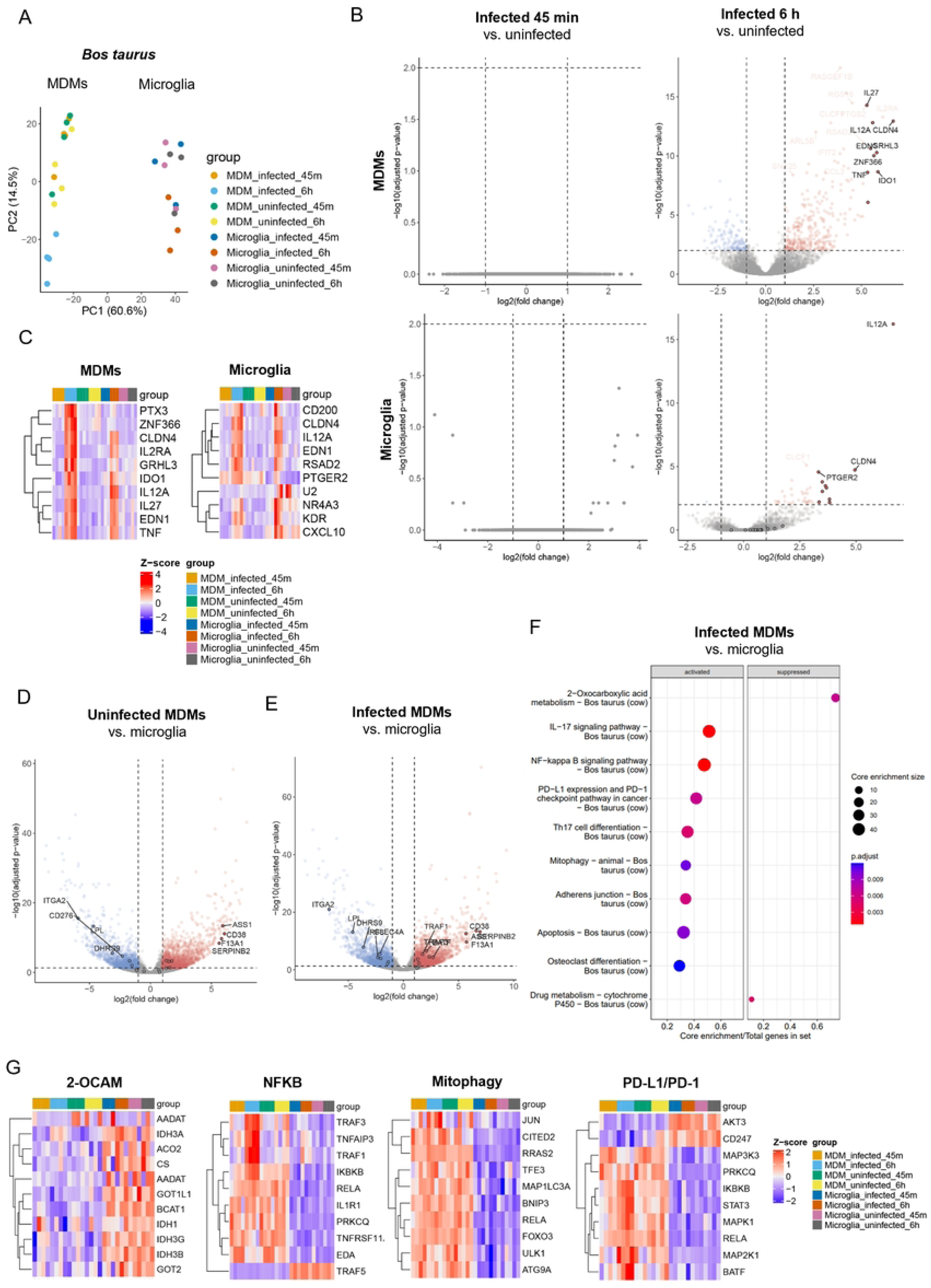
Transcriptional profiling of MDM and microglia responses to *Lm* infection. (A) Principal component analysis (PCA) of the 500 most variably expressed host genes in MDMs and microglia at 45 minutes and 6 hours pi, based on the normalized count data. Experimental groups are indicated by different colors. Each dot represents a single biological replicate from an individual animal. (B) Volcano plots comparing host transcriptomes between infected and uninfected MDMs (upper panel) and microglia (lower panel) at 45 minutes (left) and 6 hours (right) pi. Dotted lines indicate thresholds of significance (|log2FC| ≥1; padj <0.01). Transcripts that are significantly upregulated in the first condition versus the second condition are shown in red, while those that are downregulated in the first condition versus the second condition are shown in blue. (C) Heatmap of the top 10 upregulated genes in MDMs (left) and microglia (right) after 6 hours of infection as compared to uninfected cells. (D) Volcano plot comparing host transcripts between uninfected MDMs and microglia (dotted lines indicate thresholds of significance: |log2FC| ≥1; padj <0.01) at 6h pi. (E) Volcano plot comparing host transcripts between infected MDMs and microglia at 6h pi (dotted lines: |log2FC| ≥1; padj <0.01). Transcripts with significantly higher expression in MDMs are shown in red, while those with higher expression in microglia are shown in blue. (F) Top 10 significantly enriched KEGG pathways of differentially expressed host genes exclusively activated or suppressed in infected MDMs compared to infected microglia at 6 hours pi and not enriched in uninfected cells (qvalue<0.05). (G) Heatmaps of the top 10 enriched genes from the following KEGG pathways: bta01210 (2-Oxocarboxylic acid metabolism, 2-OCAM - Bos taurus), bta04064 (NF-kappa B signaling pathway - Bos taurus), bta04137 (Mitophagy - Bos taurus), bta05235 (PD-L1/PD-1 - Bos taurus). The Z-score represents scaled log-normalized counts, with genes ordered by their differential expression between infected MDMs and infected microglia at 6 hours pi.

**Figure 6.**
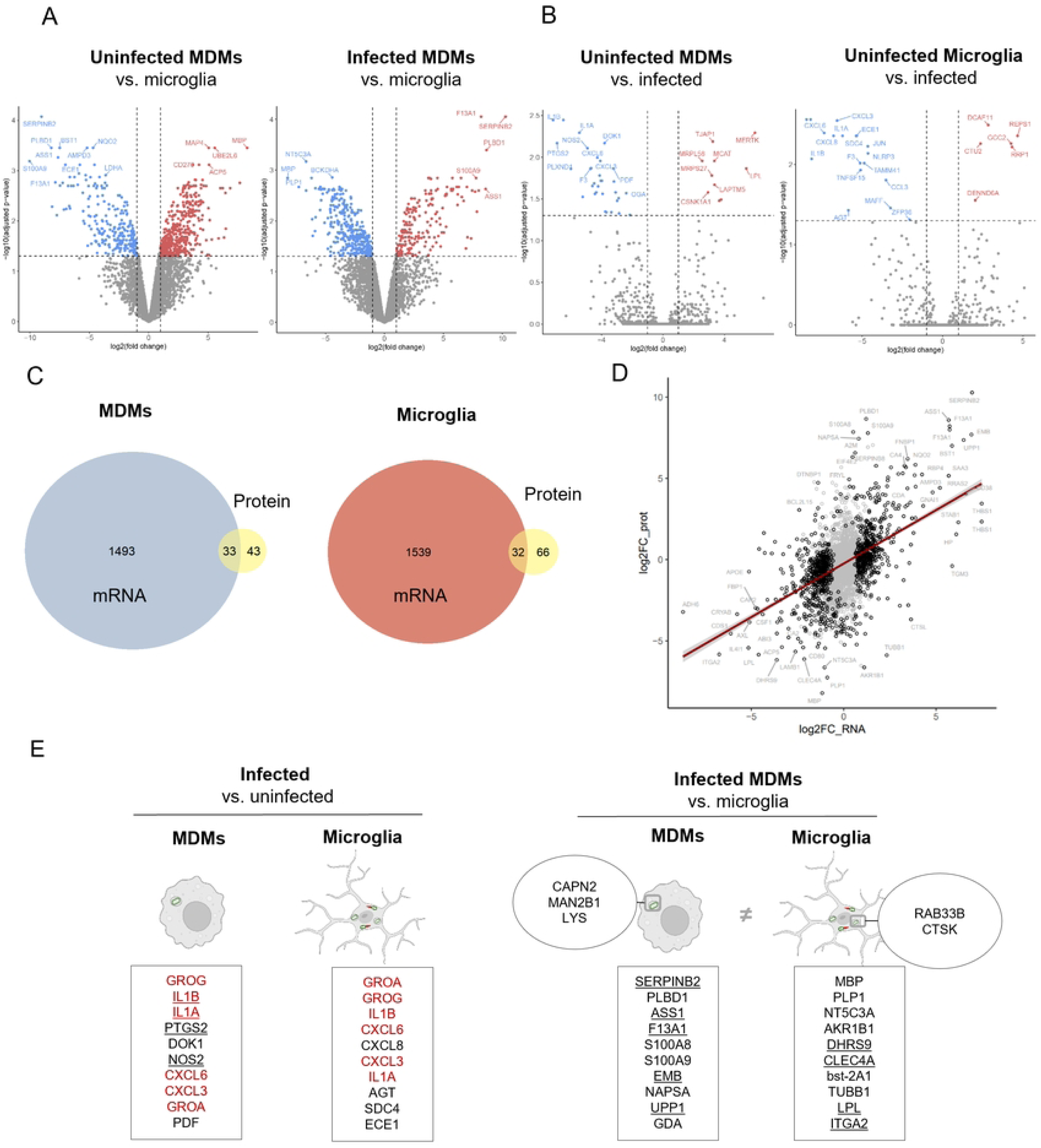
Proteomic analysis of *Lm*-infected microglia and MDMs. (A) Volcano plots comparing differentially expressed proteins (DEPs) between uninfected (left) and infected (right) microglia and MDMs (dotted lines: |log2FC| ≥1, adjusted p-value<0.05). Proteins with significantly higher expression in the first condition compared to the second condition are depicted in red, while those with lower expression in condition A versus condition B are in blue. (B) Volcano plots comparing DEPs between uninfected (red) and infected (blue) MDMs (left) and between uninfected (red) and infected (blue) microglia (right). (C-D) Correlation between protein and transcriptome responses in microglia and MDMs. (C) Venn Diagram illustrating the overlap between differentially expressed transcripts and proteins in infected MDMs and microglia. (D) Scatter plot of the linear regression between the |log2FC|≥1 of all transcripts and proteins from infected MDMs and microglia. Black: significant in either gene expression, protein abundance or both (padj < 0.01; R2=0.2032; p-value < 2E-16); Gray: not significant in any condition. (E) Schematic view of DEPs specific to infection in microglia and MDMs (|log2FC|≥1, adjusted p-value<0.05). DEPs produced by both phagocytes are shown in red and DEPs that are additionally differentially expressed on the mRNA level (|log2FC|≥1, adjusted p-value<0.05) are underlined. DEPs in squared panes: Chemokine (C-X-C motif) ligand 3, 6 and 8 (CXCL3, CXCL6, and CXCL8), Growth-Regulated Alpha Protein (GROA), Growth-regulated protein homolog gamma (GROG), interleukin-1 α (IL1A), interleukin-1 β (IL1B), Nitric Oxide Synthase 2 (NOS2), Prostaglandin-Endoperoxide Synthase 2 (PTGS2), Docking Protein 1 (DOK1), Peptide Deformylase (PDF), Angiotensinogen (AGT), Syndecan 4 (SDC4), Endothelin Converting Enzyme 1 (ECE1). (F) Schematic view of the top 10 DEPs between infected microglia and infected MDMs at 6 hours of infection, shown within squared panes (|log2FC| ≥1, adjusted p-value<0.05). Selected vacuolar genes that were significantly changed between infected and uninfected microglia compared to MDMs at 6 hours are shown within circles. DEPs that are also differentially expressed on the mRNA level (|log2FC| ≥1, adjusted p-value<0.05) are underlined. DEPs in squared panes: SERPIN domain-containing protein B2 (SERPINB2), Phospholipase B-like 1 chain A (PLBD1), Arginosuccinate synthase (ASS1), Coagulation factor XIII A chain (F13A1), Protein S100-A8 (S100A8), Protein S100-A9 (S100A9), Embigin (EMB), Napsin A aspartic peptidase (NAPSA), Uridine phosphorylase (UPP1), Guanine deaminase (GDA), Cytochrome c oxidase copper chaperone COX17 (COX17), Interferon-induced GTP-binding protein Mx1 (MX1), Solute Carrier Family 25 Member 10 (SLC25A10), bisphosphate 3-prime-nucleotidase (BPNT1), Family With Sequence Similarity 136 Member A (FAM136A), Interferon Induced Protein 44 Like (IFI44L), H1.0 Linker Histone (H1-0), Rho GTPase Activating Protein 25 (ARHGAP25), Interleukin-18 (IL18), Interferon induced protein 44 (IFI44). Selected vacuolar genes in MDMs: calpain 2(CAPN2), mannosidase α class 2B member 1 (MAN2B1), lysozyme C (LYS). Selected vacuolar genes in microglia: Ras-related protein Rab-33B (RAB33B), cathepsin K (CTSK).

Similar to *Lm* transcriptional adaptation, host responses evolved dynamically over time. At 45 minutes pi, no major differences were observed between infected and uninfected cells, whereas pronounced divergence emerged at 6 hours pi (Fig 5B). The response was more pronounced in MDMs, with 306 upregulated genes compared to 43 in microglia (Fig 5B, S2 Table). A subset of 35 upregulated genes was shared, highlighting a conserved macrophage response to *Lm*. Among the top ten induced transcripts in both populations were *IL12A*, *CLDN4,* and *EDN1* (Fig 5C). Upregulation of *IL12A* and *EDN1* is consistent with their roles in antibacterial defense, Th1 polarization and cytokine release ^85–89^. In contrast, induction of CLDN4, which encodes a tight-junction protein primarily characterized in epithelial cells ^90,91^, may indicate a previously unrecognized role in phagocyte responses to infection. Most other top transcripts encoded immunoregulatory factors upregulated in both populations (*IL2RA, IDO1, IL27, TNF, RSAD2, NR4A3, PTGER2, KDR, CXCL10*). IDO-1-mediated tryptophan depletion restricts intraphagosomal replication in *Lm-*infected dendritic cells and has been implicated in the control of other bacterial pathogens, including *Bartonella henselae* and *Coxiella burnetii* ^92–94^. *PTX3* and *ZNF366* were selectively induced in MDMs, indicating distinct functional programs in blood-borne versus tissue-resident macrophages. *PTX*3 encodes a long pentraxin family protein that enhances pathogen opsonization and clearance ^95,96^, while *ZNF*366 upregulation is indicative of early monocyte differentiation toward dendritic cells ^97^. Among transcripts associated with vacuolar trafficking, infected and uninfected microglia upregulated *GILT/IFI30*, though protein levels remained unchanged. GILT/IFI30 is required for LLO activation and phagosomal escape in mice ^98^ and is associated with inflammatory microglia in Alzheimer’s disease ^99^. GSEA of KEGG pathways showed substantial overlap between infected microglia and MDMs at 6 hours pi (S7G-H Fig). Both phagocytes enriched inflammatory pathways, including NF-kappa B (NFKB), NOD, TNF, JAK-STAT signaling and cytokine signaling (S7G-H Fig), confirming a conserved macrophage defense response. Nonetheless, enrichment patterns diverged between the two subsets (Fig 5F). Infected MDMs suppressed 2-oxocarboxylic acid metabolism (bta00190) and drug metabolism pathways (bta00982) (Fig 5 E), consistent with a metabolic shift from oxidative metabolism towards glycolysis, a feature of M1 polarization ^100,101^.

MDMs further activated apoptosis (bta04210), mitophagy (bta04137), and inflammatory signaling pathways such as PD-L1/PD-1 (bta05235), IL-17 (bta04657), Th17 differentiation (bta04657), and NFKB pathway (bta04064) compared to microglia (Fig 5E). Within the NFKB pathway, infected MDMs expressed elevated levels of *NFKB1*, *TRAF1*, *TRAF3* and *TNFAIP3*, while the basic leucine zipper ATF-like transcription factor (*BATF*) was upregulated in the PD-L1/PD-1 pathway (Fig 5F). *TRAF1* and *TRAF3* act as TNF receptor adaptors modulating NFKB signaling ^102–104^, and *BATF* regulates IL-4-dependent expression in T follicular helper cells ^105,106^. These pathway signatures are compatible with a robust M1-like polarized pro-inflammatory macrophage state in MDMs.

In summary, KEGG enrichment suggests that infected MDMs shift toward glycolysis and engage in apoptotic, mitophagy and adaptive immunity-linked programs, reflecting inflammasome activation, increased cellular stress, and M1-like polarization. In contrast, microglia maintained oxidative and detoxification pathways and limited adaptive immunity activation, consistent with their homeostatic tissue-resident role.

To complement our RNA-seq analysis, we performed quantitative untargeted proteomics using Liquid Chromatography-Mass Spectrometry (LC-MS/MS) on infected bovine MDMs and microglia at 6h (Fig 1B). In line with the transcriptomic data, proteomic differences were more pronounced between cell types (Fig 6A) than between infection states (Fig 6B). Uninfected MDMs and microglia exhibited 466 differentially expressed proteins (DEPs), with 286 upregulated in microglia and 180 in MDMs (Fig 6A). Following infection, 401 DEPs were detected (Fig 6A), with 255 upregulated in microglia and 146 in MDMs (S2 Table).

KEGG pathway analysis identified metabolic pathways (bta01100) as the major axis of divergence between infected microglia and MDMs (Suppl. Table 4), highlighting metabolic rewiring as a key determinant of macrophage activation and pathogen control ^107^. Enriched pathways including glycolysis/gluconeogenesis, carbon metabolism, the pentose phosphate pathway, the HIF-1 signaling pathway, and glutathione metabolism (bta00010, bta01200, bta00030, bta04066, bta00480) are consistent with a shift towards glycolytic proinflammatory M1-like macrophages ^101^.

Overlap between differentially expressed transcripts and proteins was limited (Fig 6C-D), likely reflecting differences in methodological sensitivity, transcript versus protein stability, post-transcriptional regulation, or different MOIs ^108,109^. Nonetheless, transcript-protein log2FC values were strongly correlated at 6 hours pi (Suppl. Table 1, Spearman correlation coefficient r=0.8696, two tailed p<0.0001).

Among the DEPs, 17 proteins upregulated in microglia and 27 in MDMs were linked to inflammation (S1, S4 Tables). Both cell types upregulated chemotactic factors such Spearman, CXCL3/GROG, CXCL6, IL1A and IL1B (Fig 6B), in agreement with transcript upregulation for selected candidates (Fig 6E). Infected MDM expressed higher levels of SERPINB2, arginosuccinate synthase 1 (ASS1), lactate dehydrogenase A (LDHA), embigin EMB, CD38, and F13A1 at both protein and mRNA levels, while S100A8/9 and phospholipase PLBD1 were upregulated only at the protein level (Fig 6E, S1 Table). MDMs also expressed markers of classical (M1-like) activation, including CD38 and TREM1, and glycolytic enzymes (LDHA, PFKL, PKM, PGK1, and GAPDH) as well as inflammatory markers (SQSTM, GPI, PGD, G6PD, and CAS9) (Suppl. Table 1, ^101^.

In contrast, infected microglia upregulated C-type lectin domain family 4 member A (CLEC4A), MERTK and the lipoprotein lipase (LPL), linked to regulation of adaptive immunity (Suppl. Table 1) and homeostatic M2 signature ^110–114^. However, they also strongly activated proinflammatory markers involved in type I interferon signaling, with elevated interferon regulatory factor 8 (IRF8) and multiple interferon-stimulated genes, including IL18, IFI44L, IFIT2, IFIT3, IFITM1, MX dynamin-like GTPase 1 (MX1), and ISG15, consistent with cytosolic pathogen recognition (Suppl.Table.1).

Within the vacuolar-associated proteins, microglia selectively enriched RAB33B, cathepsin K (CTSK) (both transcript and protein), and RAB38 (protein only), suggesting a program of phagolysosomal degradation and tissue-compatible clearance rather than the inflammatory effector profile observed in MDMs (Fig 6E; S1 Table). Conversely, MDMs exhibited higher expression of calpain 2 (CAPN2), mannosidase α class 2B member 1 (MAN2B1) (Fig 6E; S1 Table; S6 A-C Fig), matrix metallopeptidase 12 (MMP12) and cytochrome b-245 alpha chain (CYBA) (S1 Table). MAN2B1 is involved in glycoprotein catabolism, possibly suggesting increased lysosomal activity in MDMs. Metallopeptidases have been linked to reduced LLO activity and reduced escape of *Lm* in neutrophils ^115,116^. CAPN2 and CYBA promote ROS-driven antimicrobial activity typically associated with M1 activation, while MMP12 and MAN2B1 are rather linked to tissue remodeling or M2-like infiltration, together indicating a mixed vacuolar activation profile.

Notably, MDM expressed markedly higher levels of lysozyme C (LYS), over two orders of magnitude greater than microglia. Consistent with its bacteriolytic role, *Lm* was more frequently found in association with lysozyme in MDMs (S8B, C Fig). As lysozyme-positive macrophages are found within *Lm* microabscesses *in vivo* ^117^, we tested whether it contributes to the induction of VBNC *Lm.* Lysozyme treatment of extracellular bacteria inhibited bacterial growth only at high concentrations (3 mg/mL) (S7D-F Fig), in line with reported innate *Lm* resistance ^118,119^. BacLight assays showed that growth reduction reflected bacterial killing rather than VBNC induction, indicating that lysozyme alone is insufficient to induce VBNC states.

Together, these data indicate that while MDMs and microglia share core inflammatory pathways, they diverge in functional programs upon *Lm* infection. MDMs predominantly upregulate metabolic factors and antimicrobial effectors consistent with classical (M1-like) pro-inflammatory polarization, while microglia engage programs centered on phagolysosomal processing, lipid metabolism, and type I interferon signaling. This reflects a microglia-specific strategy that prioritizes immune regulation and intracytosolic pathogen sensing over broad inflammatory effector activity.

## Discussion

*Lm* is a major cause of CNS infection in humans and animals ^19,120,121^, yet the cellular and molecular mechanisms underlying neurolisteriosis remain incompletely understood. Microglia, the CNS-resident macrophages, and infiltrating MDMs play distinct roles during *Lm* infection ^24,84^. More broadly, growing evidence indicates that macrophage ontogeny shapes infection outcomes with *Lm* and other intracellular pathogens ^27,33,122,123^. However, how *Lm* adapts and exploits these ontogenetically distinct macrophage subsets remains unclear. Using dual RNA-seq paired with host proteomics, we provide the first integrated, genome-wide snapshot of macrophage subtype-specific responses to *Lm* infection and the corresponding bacterial transcriptional programs that allow *<u>Lm</u>* to survive and differentially thrive within distinct intracellular niches. Our findings reveal the biological principle that the intracellular fate of *Lm* is not determined by canonical virulence factors alone but emerges from the metabolic and immunological landscape created by macrophage ontogeny. Tissue-resident microglia and infiltrating MDM share core antimicrobial programs but diverge in nutrient provisioning, metabolic pathways and stress-imposing capacity, which in turn shape whether *Lm* adopts a replicative cytosolic lifestyle or an intravacuolar dormant, stress-tolerant phenotype. Metabolic rewiring is also a key feature of host-pathogen interactions in macrophages infected with *Leishmania infantum* and *Salmonella typhimurium* ^124–127^. On the bacterial side, differential expressions of canonical virulence genes did not account for the divergent behaviors of *Lm* in microglia and MDMs^2,52,128,129^, though shaped by the specific macrophage lineage.

In MDMs, the intracellular environment triggered downregulation of OXPHOS, PTS, fatty acid biosynthesis and metabolism, and carbon and pyrimidine metabolism (Fig 7), features consistent with a dormant-like, stress-adapted phenotype suited to survival within a hostile vacuole. Similar metabolic reprogramming support persistence in other intracellular pathogens, including *Mycobacterium*, *Salmonella*, *Francisella* and *Cryptococcus* ^130–133^. Importantly, we show that *Lm* strongly induces non-coding sRNA-mediated metabolic control, notably T-box riboswitches, which act as rapid regulators in amino acid scarcity, revealing a post-transcriptional layer of adaptation to nutrients stress ^46,75,134^. In contrast, *Lm* in microglia rapidly adopted a metabolically active, cytosolic growth program characterized by upregulation of genes involved in glycolysis, fatty acid metabolism and nucleotide salvage, enabling sustained replication (Fig 7). This mirrors strategies used by *M. marinum* during intracytosolic growth in murine BV2 microglial cells and amoeba ^131,135^, and by *M. tuberculosis* ^136–138^, which rely on fatty acids and triacylglycerols.

**Figure 7.**
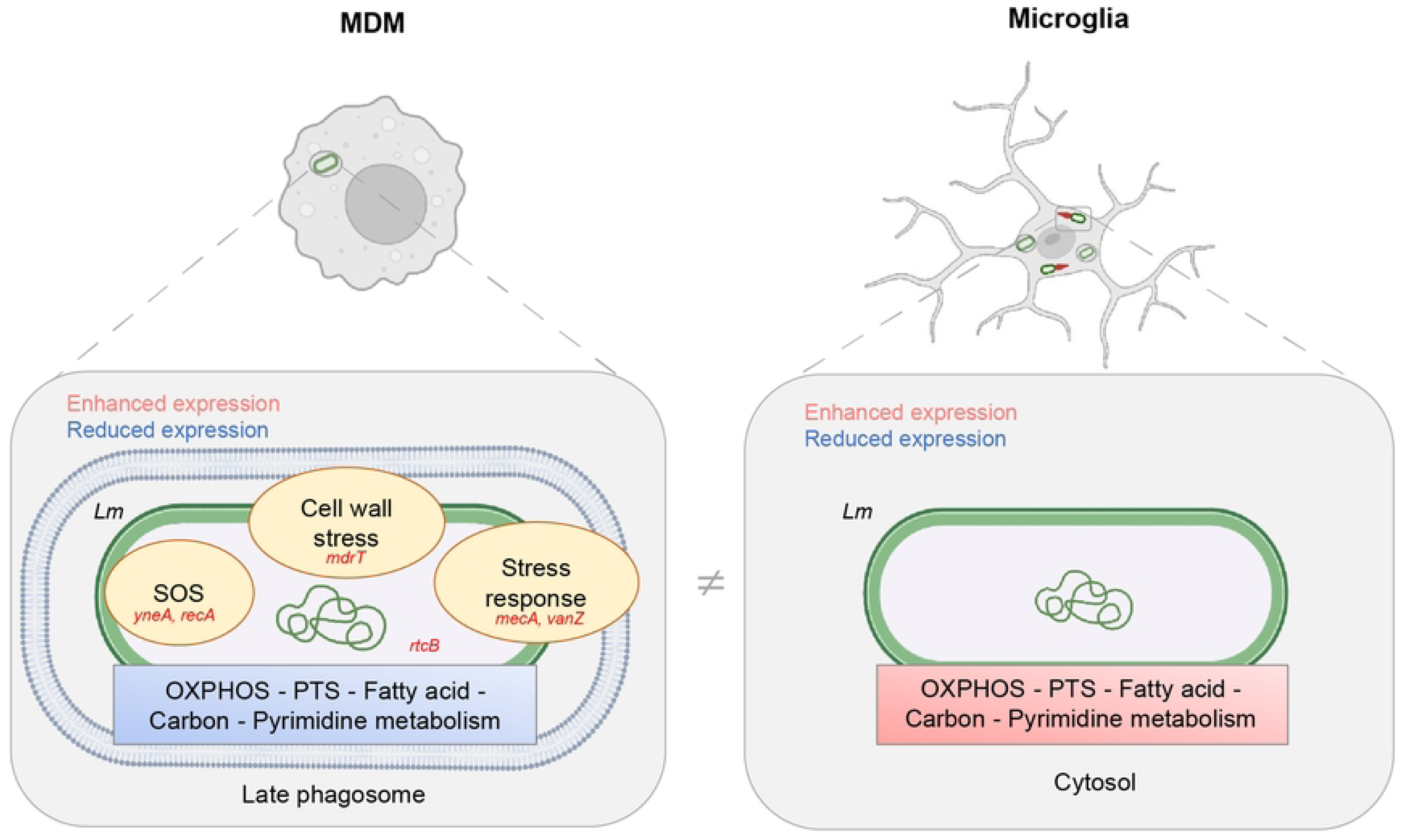
Niche specific gene expression of *Lm* in MDMs and microglia. In MDMs, *Lm* downregulates fatty acid, pyrimidine, and carbon metabolism. Moreover, the intravacuolar environment evokes a bacterial stress response and SOS response to restore DNA integrity. In microglia, the favorable intracytosolic environment results in upregulation of gene expression programs supporting oxidative and carbohydrate metabolism and host nucleotide salvage for DNA replication.

*Lm* also activated complex stress and SOS responses in MDMs. The induction of *lexA, recA* and *yneA* is consistent with a state of dormancy previously described in bovine MDMs ^33^ and resembles stress signatures in extracellular VBNC *Lm* states ^139,140^. Additional upregulated genes included multidrug transporters (*mdrT*, *mdrL)*, and antibiotic resistance determinants (*mecA*, *vanZ)* (Fig 7), classical markers of cell wall stress and antibiotic resistance in other Gram-positive bacteria ^141–144^. These findings underscore that macrophage-imposed stress may inadvertently promote the emergence of dormancy-associated phenotypes with reduced antibiotic susceptibility. The strong induction of *rtcB*, a tRNA ligase involved in RNA repair first described in *E. coli* ^145^, further supports the activation of complex stress-compensatory pathways. Collectively, these transcriptional signatures resemble dormancy programs described across other persistent bacterial pathogens ^83,131,146,147^. Functionally, we identify *recA* and *rtcB* as contributors to intracellular persistence in a cell-type specific manner. Deletion of either gene increased VBNC formation in microglia, whereas in MDM only *recA* deletion exacerbated the VBNC-like phenotype in MDMs, suggesting compensation of *rtcB* loss by redundant RNA repair pathways in the latter. The intravacuolar restriction observed for the *recA* and *rtcB* deletion mutants in microglia at 24 hours likely reflects reduced bacterial fitness under phagosome stress, impairing the critical transition to the cytosolic niche. One possible explanation is that RecA and RtcB are transiently required for bacterial survival within microglial phagosomes prior to escape. Such a requirement may not be captured by transcriptomic analyses performed as early as 45 min post-infection due to limited sensitivity or because these processes are regulated at the post-transcriptional or post-translational level.

On the host side, both resident microglia and infiltrating MDM mounted a robust IL-1-driven inflammatory cascade, producing IL1B, IL1A, CXCL1, CXCL3, and CXCL6, indicative of neutrophil-recruiting pro-inflammatory responses. At a late stage, however, striking divergences became evident, reflecting ontogeny-driven host immune programming. MDMs adopted a glycolytic, pro-inflammatory M1-like profile ^148,149^ and upregulated phagolysosomal effectors such as MAN2B1, LYS and NOS2, generating a hostile intravacuolar environment that restricts cytosolic escape and imposes metabolic and oxidative pressure. In contrast, microglia showed a hybrid inflammatory and reparative phenotype. They strongly induced type I interferon (IFN-I)-related genes (IRF8, IFI44L, IFIT2, IFIT3, and IFITM1, Suppl. Table 1), reflecting IFN-I activation during cytosolic *Lm* infection ^150^ ^150,151^ along with PTGS1, supporting an M1-like proinflammatory response. Yet, they simultaneously expressed M2-like markers associated with homeostatic or inflammation-resolving states (LPL and MERTK). These features reinforce the concept that the binary M1/M2 paradigm insufficiently captures the functional and metabolic plasticity of macrophages during infections^152^.

The delayed host responses (6 hours) compared to the timing of vacuolar escape reported in mice (30 min) ^51^ suggest that the cell-type specific fate of *Lm* is not dictated by any single activated host pathway but instead emerges from the coupling of bacterial survival strategies with the metabolic and immunological program inherent to the macrophage lineage, further supporting the principle that macrophage ontogeny shapes pathogen fate ^41^. Both macrophage types exert antimicrobial pressure, yet their intracellular environments lead to opposing outcomes: MDMs drive *Lm* into a stressed, dormant-like state, whereas microglia serve as amplification niches for *Lm* within the CNS. Unlike MDMs, microglia maintain a metabolically active cytosol and imposes comparatively low intracellular stress, permitting *Lm* to complete vacuolar escape, replicate, and spread. This suggests that CNS susceptibility to *Lm* may arise in part from the presence of a resident macrophage population inherently permissive to intracellular growth, a concept that may extend to other neurotropic pathogens. In conclusion, our work shows that the pathogenesis of neurolisteriosis arises from the interplay between bacterial virulence strategies and macrophage lineage-specific host programs, rather than from canonical virulence factors alone. *Lm* flexibly adopts persistence and growth programs according to the intracellular niche, thereby maximizing its survival within the host. Our study further identifies *recA* and *rtcB* as contributors to intracellular persistence, with distinct functional relevance in microglia and MDMs. These findings underscore macrophage ontogeny as a critical determinant of infection phenotype, advance the conceptual framework of host-imposed metabolic landscapes as drivers of pathogen adaptation, and highlight the importance to account for macrophage diversity when developing diagnostic and therapeutic strategies for CNS infections.

## Methods

### Isolation and in vitro culture of primary cells and cell lines

Primary bovine microglia and MDMs were isolated and cultured as previously described ^1^. Briefly, brain cortex tissue and blood were obtained from cattle younger than one year of age, slaughtered at a local abattoir for human consumption. Microglia were isolated by mechanical dissociation followed by centrifugation on a 37% Percoll gradient. Monocytes were isolated from peripheral blood mononuclear cells (PBMC) using CD14+ magnetic-activated cell sorting (Milteny Biotec) and differentiated into MDMs. Both microglia and MDMs were maintained in Dulbecco’s modified Eagle’s medium (DMEM; Invitrogen, Gibco) supplemented with 10% heat-inactivated fetal bovine serum (FBS) (Capricorn Scientific), 1% penicillin/streptomycin (P/S) (Invitrogen, Gibco), and recombinant macrophage colony-stimulating factor (M-CSF; 25 ng/mL Kingfisher, Biotech, Inc.). Cells were used for infection experiments after seven days of culture. The human microglial cell line ^153^ maintained under the same conditions as bovine primary cells (DMEM, 10% heat-inactivated FBS, P/S).

### Bacterial strains and cultures

The *Lm* strain JF5203 (lineage I, clonal complex 1, sequence type 1, https://www.ncbi.nLm.nih.gov/nuccore/NZ_LT985474.1) was used as the parental wild-type strain (wt-*Lm*). This strain was originally isolated from a case of bovine rhombencephalitis, and GFP-expressing JF5203 *Lm* were generated in previous studies ^21,120,121^. Deletion mutants (Δ*mecA*, Δ*vanZ*, Δ*mdrT*, Δ*yneA*, Δ*rtcB,* Δ*recA*) were generated by homologous recombination using the temperature-sensitive plasmid pMAD, following the protocol of Boneca et al. ^154^ with adaptations. Synthetic gene fragments containing fused flanking regions of the target genes (Table 1) were synthetized and cloned into pMAD by Twist Bioscience (Suppl. Table 7).

Plasmids were introduced into electrocompetent wt-*Lm* by electroporation in a 0.1-cm cuvette using a GenePuls apparatus (Bio-Rad, Hercules, CA, USA) with settings of 25 μF, 200 Ω, and 1.25 kV ^155^. Transformants were selected on brain heart infusion (BHI) agar plates containing 5 μg/ml erythromycin (Sigma) and 50 μg/ml X-Gal (5-bromo-4-chloro-3-indolyl-β-d-galactopyranoside) at 30°C for 2 days. A single blue colony was grown in BHI broth with 5 μg/ml erythromycin at 42°C for 2 days, followed by serial dilutions plated onto BHI agar with erythromycin (5 μg/ml) and X-Gal (50 μg/ml) at 42°C for 2 days. A single blue colony was transferred into BHI broth and grown at 30°C for 6 hours, followed by subculturing in fresh BHI broth at 42°C for 18 h. Serial dilutions were plated onto BHI agar with X-Gal and grown at 42°C for 2 days. White colonies, indicative of plasmid excision, were screened for erythromycin sensitivity and verified by PCR for loss of the pMAD backbone and the targeted gene deletion as well as the presence of the *Lm hly* gene (S5 Fig).

All mutants and the parental strain underwent PacBio whole-genome sequencing and sequence analysis using SnpEff (https://pcingola.github.io/SnpEff) ^156^ to confirm accurate gene deletions and screen for off-target mutations (S5 Fig, S7 Table).

### Axenic growth curves and lysozyme susceptibility

Axenic curves were generated as previously reported ^157,158^. Stationary-phase overnight BHI cultures of single colonies of wt-*Lm* and deletion mutants were diluted 1:40 (on average OD600=0.025) into fresh BHI and incubated at 37°C in 96 well plates. OD600 was recorded every 30 minutes for 12 hours (three independent experiments in technical triplicates). For lysozyme susceptibility assays, wt-*Lm* was grown to mid-log phase and exposed to serial dilutions of hen egg white lysozyme (Sigma Aldrich) in PBS. OD600 was measured hourly for 6 hours and normalized to untreated controls.

### Infection assays

Primary microglia, MDMs, and the HMC-3 cell line were infected in gentamicin protection assays. Eighteen hours prior to infection, the culture medium was replaced with DMEM supplemented with 10% FBS and recombinant M-CSF (25 ng/mL) without antibiotics. One hour before infection, the medium was changed to DMEM without FBS. Overnight bacterial cultures of wt-*Lm* and deletion mutants were diluted in PBS to OD600=0.8 and added at a multiplicity of infection (MOI) of 5 (phenotypic assays), 10 (proteomics), or 20 (dual RNA-seq). Higher MOI was used to improve bacterial transcript/protein recovery ^46^ without altering infection dynamics. To synchronize infection, plates were centrifuged at 250g for 2 minutes. After 30 minutes, cells were carefully washed with PBS and maintained in DMEM supplemented with 10% FBS and 50 μg/ml gentamicin (Sigma-Aldrich) to eliminate extracellular bacteria. Infection readouts included CFU quantification, bulk dual RNA-sequencing, proteomics, and confocal microscopy at defined time points.

### Colony Forming Units (CFU) quantification

At 2, 6, and 24 hours after infection, cells were washed twice with PBS and lysed with 0,5% TritonX-100 (Sigma Aldrich). Serial ten-fold dilutions (1:1 to 1:10,000) were plated on BHI-agar for CFU enumeration. Counts were normalized to the initial inoculum. Unless indicated otherwise, data represent at least three independent biological replicates performed in technical triplicates.

### Bulk dual RNA sequencing RNA isolation and sequencing

RNA was extracted from infected primary bovine microglia (n=3) and MDMs (n=4), as well as matched mock-infected microglia and MDMs controls the same animals, and *Lm* grown in FBS (n=3) to model extracellular bacteria in blood. Microglia and MDMs were seeded at 500,000 and 3 million cells per well in 24-well plates and 6-well plates, respectively, and infected with wt-*Lm*-GFP (MOI=20) in a gentamicin protection assay. RNA was extracted at 45 minutes and 6 hours pi. Bacterial RNA was obtained from *Lm* cultured in FBS (1:10) for 6 hours. Host and bacterial RNA were co-released using enzymatic permeabilization (20mg/ml lysozyme, Tris HCl 20 mM, EDTA 2mM, 1.2% Triton X-100) for 20 minutes at 37°C ^159^. Total RNA was extracted using TRI Reagent® and the Direct-zol Miniprep kit (Zymo, R2050). RNA quality and quantity were assessed using an Agilent 2100 Bioanalyzer (Agilent Technologies) and a Qubit 2.0 Fluorometer (Life Technologies), respectively. The mean RNA integrity number (RIN) was 8.4, and the minimum was 6. After rRNA depletion (Illumina Ribo-Zero plus rRNA Depletion Kit), total paired-end mRNA libraries were prepared using the CORALL Total RNA-Seq Kit (Lexogen) with Unique Molecular Identifiers (UMI) for deduplication. Sequencing was performed on Illumina NovaSeq 6000 SPFlow cell platform with a sequencing depth of 48 million read pairs per sample.

### Bioinformatics analysis

RNA-seq quality was assessed using Fastqc v.0.11.9 (Andrews S 2010, https://www.bioinformatics.babraham.ac.uk/projects/fastqc/) and RSeQC v.4.0.0 ^161^. Reads were aligned to a concatenated Bos taurus (ARS-UCD1.2; GCF_002263795.1) and *Lm* (JF5203_1; GCF_900275645.1) genome using HiSat2 v.2.2.1 ^162^. Read counts were obtained with FeatureCounts v.2.0.1 ^163^, separately analyzing host and bacterial transcripts based on their respective genome annotations. All further analyses were run in R version 4.2.1 (2022-06-23) (R Core Team. 2022 https://www.r-project.org). Differential gene expression (DGE) analysis was performed using the Bioconductor package DESeq2 v1.38.3 ^164^, with significance thresholds of |log2FC| ≥ 1 (FC= fold changes) and padj < 0.01 for host genes and < 0.05 for *Lm* genes. Gene set enrichment analysis (GSEA) ^165^ was run in ClusterProfiler v4.7.1 ^166^ using KEGG ^167^ and MSigDb ^168^gene sets.*Lm* gene ontology (GO) analysis was conducted in STRING using the EGD-e annotation, differentially expressed genes between conditions and False Discovery Rate (FDR) < 0.05. The EGD-e annotation was also used to plot STRING networks. Small RNAs were analyzed by combining and indexing the cow genome with the EGD-e genome for mapping, re-aligning the reads and counted alignments to the EGD-e ORF and small RNAs ^75^, followed by differential gene expression analysis of small RNAs.

### Proteomics analysis

Quantitative untargeted liquid chromatography-tandem mass spectrometry (LC-MS/MS) was performed on infected microglia (n=3) and MDMs (n=3), and matched mock-infected controls (n=3). Cells were infected at MOI=10 (6*106 CFU/well) for 6 hours, harvested with trypsin, and resuspended in DMEM with FBS. Cells were washed twice to remove traces of FBS and lysed in 8M urea, 100mM Tris/HCl (pH=8), containing EDTA-free protease cocktail inhibitor (Roche, Rotkreuz, Switzerland). Lysates were snap-frozen in liquid nitrogen and stored at -80°C before further processing and analysis, as previously described ^169^. Mass spectrometry data were analyzed for comparison and quantification against a concatenated database consisting of all Uniprotkb Bos taurus sequences (version 2022 12) ^170^ and common contaminants ^171^ using fragpipe version 19.1 ^172^. Search parameters included precursor and fragment mass tolerances of ±10 ppm and 0.4 Da, full trypsin cleavage specificity allowing for up to three missed cleavages, fixed carbamidomethyl modification of cysteines, variable N-terminal acetylation and oxidation of methionine. Protein intensities were normalized using the MaxLFQ method, and differential expression between groups was analyzed with the Empirical Bayes test (FDR-adjusted p < 0.05, |log2FC| ≥ 1). Kyoto Encyclopedia of Genes and Genomes (KEGG) analysis of differentially expressed proteins (DEPs) was performed using Database for Annotation, Visualization, and Integrated Discovery (DAVID). For integrative analysis, transcriptomic and proteomic datasets (from 6h pi) were combined, and regression analysis was applied to assess correlations between DEPs and DEGs ^173^. Spearman correlation between mRNA and protein log2FC was computed using GraphPad Prism (version 10.0.0, GraphPad Software, San 324 Diego, California USA, www.graphpad.com).

### Bacterial viability assay

MDMs and microglia were seeded on poly-L-lysine-coated (P6282, Sigma) glass coverslips or 96-well plates (IBIDI®, GMBH) at densities of 250’000 and 90’000 cells per well, respectively. Intracellular bacterial viability was assessed at 6 and 24 h pi using the LIVE/DEAD BacLight Bacterial Viability Kit (L7012, Life Technologies) in a gentamicin protection assay as previously described ^33,36^. Cells were washed twice in 0.1 M 3-(N-morpholino) propanesulfonic acid (MOPS, pH 7.2) containing 1 mM MgCl2 (MOPS/MgCl2) and incubated for 15 minutes at room temperature in the dark with Live/Dead Staining Solution (1.6 μM SYTO9, 20 μM propidium iodide, and 0.1% saponin) in MOPS/MgCl2. Imaging was performed within 30 minutes using an Olympus FV3000 confocal laser scanning microscope. Ten random fields of view (FOVs) were acquired per coverslip and processed and analyzed with Fiji (imagej.net).

### Immunofluorescence (IF)

Infected cells were washed with PBS and fixed in 4% paraformaldehyde (PFA) for 30 minutes at 37°C, followed by PBS-0,1% Tween (PBS-T) washes and permeabilization with 0,2% Triton-X100 in PBS for 5 minutes. Primary antibodies against lysozyme (A0099, polyclonal rabbit anti-human; Dako, 1:200) or LAMP1 (ab24170, abcam, 1:100) were applied in PBS-T with 10% normal goat serum (NGS) for four hours or one hour at room temperature (RT), respectively. After PBS-T washes, cells were incubated with secondary antibody (AlexaFluor 555 IgG goat anti-rabbit; Life Technologies, 1:500), DAPI (Thermo Fisher Scientific, 1:1000), and phalloidin DyLight 633 (Invitrogen, 1:300) in PBS-T with 10% NGS for one hour in the dark at RT. For double IF experiments, Zenon-488 (Invitrogen, 1:500) labelled rabbit Listeria O antiserum (Difco, 1:200) was added after the secondary antibody incubation. Coverslips were mounted using Glycergel (Dako), and images were acquired using an Olympus FV3000 confocal laser scanning microscope. At least 10 representative FOVs per time-point were analyzed per experiment using Fiji (imagej.net). Bacteria and macrophage cells were counted using the Cell Counter plugin in Fiji. The proportions of LAMP1-associated bacteria were quantified relative to intravacuolar bacteria (lacking actin polymerization) and actin cloud or tail-associated cytosolic bacteria to the total bacterial population.

### Statistical analysis

Statistical analysis was conducted in GraphPad Prism (version 10.3.1, GraphPad Software, San 324 Diego, California USA, www.graphpad.com). Data are presented as mean with standard error of the mean (SEM). Comparisons between groups were made using non-parametric Mann-Whitney U tests or parametric Student’s t tests, as appropriate. Linear regression analyses were performed in R-studio. Significance thresholds are indicated in figure legends.

## Acknowledgements

This research was funded by the Swiss National Science Foundation (Grant No. 310030_197879). We thank the slaughterhouse Metzgerei Holzer in Hindelbank and Metzgerschaft Berner Oberland in Thun, for providing bovine brain and blood samples. RNA sequencing and proteome analysis were conducted at the Next Generation Sequencing (NGS) platform of the Vetsuisse faculty and the Proteomics and Mass Spectrometry Core Facility (PMSCF) of the Department for BioMedical Research (DBMR) at the University of Bern, which we acknowledge. We thank Dr. Pamela Nicholson, Prof. Manfred Heller, Prof. J. Frey and Prof. M. Loessner for their insightful discussions. The authors used ChatGPT (OpenAI) to assist with language editing of the manuscript. The authors reviewed and edited the AI-generated output and take full responsibility for the content of this article.

## Data Availability

Raw proteomics datasets from infected and uninfected bovine primary microglia and MDM have been deposited in the PRIDE depository under the accession number PXD071215. Raw dual RNA-seq are available in ENA/EMBL-EBI (ArrayExpress) depository under the accession number: E-MTAB-16397. The datasets are currently under embargo and will be made publicly accessible upon acceptance of the manuscript. All processed data are provided within the manuscript and supplementary materials.

## Author Contributions

Conceptualization: Anna Oevermann; Experimental design: Margherita Polidori, Anna Oevermann; Collection of animal tissues: Margherita Polidori, Camille Monney; Conduction of experiments: Margherita Polidori, Camille Monney; Data collection and processing: Margherita Polidori, Geert Van Geest, Alba M. Neher; Data analysis: Margherita Polidori, Geert Van Geest, Remy Bruggmann, Anna Oevermann. Writing: Margherita Polidori, Anna Oevermann; Funding acquisition: Anna Oevermann; Resources: Division of Neurological Sciences, Vetsuisse Faculty, University of Bern; Supervision: Anna Oevermann.

## Supplementary Figures

**Supplementary Figure 1. Quality control plots of host read mapping and counting.** (A) Read pairs aligned to the bovine reference genome. To assign reads to a gene, read pairs were aligned to the bovine reference genome. It is expected that most reads map uniquely to a region on the genome. (B) Aligned reads that were assigned to genes. The more alignments are assigned to a gene, the higher the abundance of the mRNA. Often, many alignments cannot be assigned to a gene, due to alignment to a non-genic region or alignment to multiple genes for instance.

**Supplementary Figure 2. Overview of bacterial intracellular transcripts versus extracellular control.** (A-B) Volcano plots of bacterial transcripts that were significantly changed between MDMs or microglia and FBS (dotted line: |log2FC| ≥1; padj<0.05). Transcripts that are significantly upregulated in the first condition versus the second condition are shown in red, while those that are downregulated are shown in blue. (A) *Lm* transcripts in MDMs or microglia infected for 6 hours (right) compared to FBS (left). (B) *Lm* sRNA transcripts in infected MDMs or microglia for 6 hours (right) as compared to FBS (left). (C-D) KEGG pathways of *Lm* transcripts that are activated or suppressed within infected MDMs or microglia compared to FBS (qvalue<0.05). (C) Bacterial KEGG pathways enriched in MDMs at 6 hours pi versus FBS. (D) Bacterial KEGG pathways enriched in microglia at 6 hours versus FBS. (E-F) KEGG pathways of *Lm* transcripts that are activated or suppressed over time (qvalue<0.05). (E) Bacterial KEGG pathways enriched in infected MDMs at 6 hours pi as compared to 45 minutes. (F) Bacterial KEGG pathways enriched in infected microglia at 6 hours as compared to 45 minutes pi.

**Supplementary Figure 3. Small RNAs of *Listeria monocytogenes* in infected microglia and MDMs.** (A) PCA of the 500 most variably expressed bacterial sRNAs in extracellular bacteria grown in FBS for 6 hours and intracellular bacteria in MDMs and microglia at 45 minutes pi and 6 hours pi. Experimental groups are indicated by different colors. Each dot represents a single biological replicate from an individual animal or bacterial culture. (B-F) Volcano plots of bacterial small RNAs (sRNAs) that were significantly changed between phagocytes, over time, or with FBS (dotted line: |log2FC| ≥1; padj<0.05). Transcripts that are significantly upregulated in the first condition versus the second condition are shown in red, while those that are downregulated in the first condition versus the second condition are shown in blue. (B) Bacterial sRNAs in MDMs compared to microglia infected for 45 minutes and 6 hours after infection are shown in the left and right panel, respectively. Bacterial sRNAs that were upregulated in MDMs are shown in red, while those upregulated in microglia are shown in blue. (C) Bacterial sRNAs that were upregulated in MDMs (left panel, red) or microglia (right panel, red) after 45 minutes of infection or in FBS (blue). (D) Bacterial sRNAs in MDMs (left panel) or microglia (right panel) upregulated at 6 hours (red) compared to 45 minutes pi (blue).

Supplementary Figure 4. STRING network of protein-protein interaction from the list of *Lm* genes upregulated in phagocytes over time. (A-B) STRING networks (high confidence = 0.700) showing bacterial genes that were differentially expressed over time (6 hours vs. 45 minutes) in infected MDMs and microglia. Network nodes represent proteins, while filled nodes show a known or predicted 3D structure of a protein, and edges represent protein-protein associations. (A) STRING network of *Lm* DEGs upregulated MDMs infected for 6h compared to 45 minutes pi. (B) STRING network of *Lm* DEGs upregulated microglia infected for 6 hours compared to 45 minutes pi.

**Supplementary Figure 5. Analysis of the genome of deletion mutants.** (A) Electrophoresis gel of a PCR confirming gene deletions in the mutants. Primers were used to target the deleted genes and a conserved *Lm* gene (*hly*). From left to right: 1kb ladder, 1: deleted gene in deletion mutant, 2: positive control (expressed gene in wt-*Lm*), 3: negative control (DNase free water), 4: *hly* expression in the deletion mutant. B: Table showing the primers used in this study. C: Sequence analysis of off target mutations classified with high, moderate and low impact. Further sequence variations included a GCA→GTA substitution in CW42_RS05475 in the Δ*rtcB* mutant (predicted high impact), and a CCA →TCA substitution in CW42_RS14430 in the Δ*recA* mutant (predicted moderate impact). The Δ*vanZ* mutant contained a disruptive in-frame deletion of 18 bp in C6W42_RS05815, and a 1 bp deletion in C6W42_RS11915, while the Δ*yneA* strain harbored a 3 bp disruptive in-frame deletion in C6W42_RS01015. Given that both deletion mutants carrying sequence variants displayed phenotypes indistinguishable from the parental strain under infection conditions, we consider these secondary mutations unlikely to be relevant.

**Supplementary Figure 6. Infection dynamics of different bacterial deletion mutants.** (A) Axenic growth curve of deletion mutants (Δ*mecA*, Δ*vanZ*, Δ*mdrT*, Δ*yneA*, Δ*rtcB*, and Δ*recA*) and parental strain (wt-*Lm*). All strains were grown overnight in BHI medium at 37°C, inoculated into fresh broth at an OD600 of 0.05 and cultured at 37°C for 12 hours. OD600 was measured in intervals of 30 minutes. Results from three independent experiments in triplicates are indicated as mean, and error bars indicate the standard deviation (SD). Statistical analysis with unpaired t-student test did not reveal any significant difference in extracellular growth between deletion mutants (Δ*mecA*, Δ*vanZ*, Δ*mdrT*, Δ*yneA*, Δ*rtcB*, and Δ*recA*) and wt-*Lm*. (B-D) Gentamicin protection assays using the HMC-3 human microglia cell line and primary MDM and microglia. (B-D) CFU quantification in HMC-3 cells (wt-*Lm*, Δ*recA*, Δ*rtcB*) and primary cells (wt-*Lm*, Δ*mecA*, Δ*vanZ*, Δ*mdrT*, Δ*yneA*) were performed in four independent experiments, with three technical triplicates in MDMs and HMC-3, and duplicates in microglia. Intracellular CFUs were normalized to the inoculum, and data are shown as mean ± SEM. (B): HMC-3 cells infected with wt-*Lm* (control), Δ*rtcB*, and Δ*recA*. Mann-Whitney test comparing wt-*Lm* with Δ*recA* and Δ*rtcB* at 24h (**p<0.01). (C) MDMs (left panel) and microglia (right panel) infected with wt-*Lm* (control), Δ*mecA*, Δ*vanZ*, Δ*mdrT*, and Δ*yneA.* (D) BacLight assay showing SYTO9+ viable bacteria per cell in MDMs (left panel) and microglia (right panel) infected with wt-*Lm* (control), Δ*mecA*, Δ*vanZ*, Δ*mdrT*, and Δ*yneA*. Quantifications of bacteria in BacLight assays were obtained from three biological replicates. (E) Confocal images of MDMs and microglia infected with Δ*mecA*, Δ*vanZ*, Δ*mdrT*, and Δ*yneA* strains (panel above) for 6 and 24 hours, and (F) wt-*Lm*, Δ*recA*, or Δ*rtcB* at 6 hours pi. Live bacteria are stained in green using SYTO9, while dead bacteria in red using PI. The inset provides a higher-magnification view of the highlighted region. (G) Intracellular CFU are compared to SYTO9-stained live bacteria, measured simultaneously in three experiments in triplicates and normalized to the host cell counts of MDMs and microglia infected with wt-*Lm*, Δ*mecA*, Δ*vanZ*, Δ*mdrT*, Δ*yneA*, Δ*recA*, or Δ*rtcB* for 6 and 24 hours in a gentamicin protection assay. Intracellular CFU are compared to SYTO9-stained live bacteria, measured simultaneously in three experiments (in triplicates) and normalized to the host cell counts. Error bars indicate the standard error of the mean (SEM).

**Supplementary Figure 7**. **Different host transcriptional profiles between cell types and over time.** (A-D) Volcano plot of host transcripts that were significantly changed between cell types and in uninfected samples (dotted lines: |log2FC| ≥1; padj<0.01). (A-B) Changes in transcripts between host cell types. (A) Infected MDMs (red) versus infected microglia (blue) at 45 minutes pi. (B) Uninfected MDMs (red) and microglia (blue) at 45 minutes pi. (C-F) Changes over time in the transcriptome of the host (dotted line: |log2FC| ≥1; padj<0.01). (C) MDMs infected for 6 hours (red) versus MDMs infected for 45 minutes (blue). (D) Microglia infected for 6 hours (red) versus microglia infected for 45 minutes (blue). (E) Uninfected MDMs at 6 hours (red) versus 45 minutes (blue). (F) Uninfected microglia at 6 hours (red) versus uninfected microglia at 45 minutes (blue). (G-H) KEGG pathways of differentially expressed host transcripts that are activated or suppressed after 6 hours of infection (qvalue<0.05). (G) Top 10 KEGG pathways enriched in infected MDMs as compared to uninfected MDMs 6 hours pi. (H) Top 10 KEGG pathways enriched in infected microglia as compared to uninfected cells 6h pi. (I) Heatmap of mRNA expression of microglia and MDMs signature genes. *P2RY12, SALL1* and *APOE* mRNA levels were significantly higher in uninfected microglia compared MDMs after 45 minutes, while *F13A1, CD14, CD163, CD209* and *EMILIN2* were higher in MDMs.

**Supplementary Figure 8. MDMs display significantly higher levels of lysozyme compared to microglia.** (A) Lysozyme C (protein ID = P80189) protein levels expressed as normalized protein intensities of infected and uninfected microglia and MDMs protein-imputed iMaxLFQ measurements. Infected and uninfected MDMs display significantly higher levels of lysozyme as compared to microglia (Error bars indicate SEM. Unpaired t-test, *p <0.05). (B) Immunofluorescence images of MDMs and microglia infected for 6h at MOI=20 (OD600=0.8) in a gentamicin protection assay. The inset shows the association between lysozyme and wt-*Lm*. Red: phalloidin; Magenta: Lysozyme; Green: wt-*Lm*-GFP; Cyan: DAPI. (C) Percentage of wt-*Lm* associated with lysozyme in MDMs and microglia. Bacteria were more frequently associated to lysozyme in MDMs than in microglia (Error bars indicate SEM. Unpaired t-test, * p < 0.05). (D) Axenic growth curves of wt-*Lm* grown in BHI broth (n=3) and treated with serial titrations of lysozyme (0.3 µg/ml, 3 µg/ml, 30 µg/ml, 300 µg/ml, 3000 µg/ml). Concentration of 0.3 µg/ml was estimated on the base of relative abundance measurements from proteome data. (E) CFU in triplicates from wt-*Lm* grown in BHI broth (n=2) and treated with 300 µg/ml or 3000 µg/ml of lysozyme. (F) Confocal image of bacteria treated with 1 mg/ml or 2 mg/ml of lysozyme for 2 and 6 hours and stained with Baclight LIVE/DEAD. Live bacteria are stained in green using SYTO9, while dead bacteria in red using PI. Treated bacteria form large aggregates as early as 2 hours and are overall fewer as compared to wt-*Lm*. Aggregates are more prominent at higher lysozyme concentration of 2mg/ml. Lysozyme is reported as LZ in the legends.

## Supplementary Tables

**Supplementary Table 1.** List of total host mRNAs expressed in infected microglia and MDMs at 6 hours pi, and infected MDMs or microglia versus uninfected phagocytes at 6h pi. List of total proteins across all conditions. Log2FC of combined DEGs and DEPs in infected MDMs versus infected microglia at 6 hours pi.

**Supplementary Table 2.** Overview of *Lm* and host DEGs and DEPs across all conditions.

**Supplementary Table 3.** Expression of selected *Lm* key intracellular virulence genes and stress response genes in MDMs and microglia across the different groups and conditions.

**Supplementary Table 4.** KEGG pathways from DEPs between infected MDMs and microglia after 6 hours of infection.

**Supplementary Table 5.** Sequences used to generate deletion mutants and primers used in this study.

**Supplementary Table 6.** Differentially expressed small RNAs across all conditions

**Supplementary Table 7.** Overview of the generation of the deletion mutants and SnpEff analysis.

## References

1. Beek, D. van de & Brouwer, M. C. Neurological infections in 2023: surveillance and prevention. Lancet Neurol. 23, 30–32 (2024).

2. Radoshevich, L. & Cossart, P. Listeria monocytogenes: towards a complete picture of its physiology and pathogenesis. Nat. Rev. Microbiol. 2017 161 16, 32–46 (2017).

3. Murray, E. G. D., Webb, R. A. & Swann, M. B. R. A disease of rabbits characterised by a large mononuclear leucocytosis, caused by a hitherto undescribed bacillus Bacterium monocytogenes (n.sp.). J. Pathol. Bacteriol. 29, 407–439 (1926).

4. Brouwer, M. C., Beek, D. van de, Heckenberg, S. G. B., Spanjaard, L. & Gans, J. de. Community-Acquired Listeria monocytogenes Meningitis in Adults. Clin. Infect. Dis. 43, 1233–1238 (2006).

5. Durand, M. L. et al. Acute bacterial meningitis in adults. A review of 493 episodes. *N. Engl. J. Med.* **328**, 21–28 (1993).

6. Schuchat, A. et al. Bacterial meningitis in the United States in 1995. Active Surveillance Team. N. Engl. J. Med. 337, 970–976 (1997).

7. Sigurdardóttir, B., Björnsson, O. M., Jónsdóttir, K. E., Erlendsdóttir, H. & Gudmundsson, S. Acute bacterial meningitis in adults. A 20-year overview. Arch. Intern. Med. 157, 425–430 (1997).

8. Gaballa, A., Guariglia-Oropeza, V., Wiedmann, M. & Boor, K. J. Cross Talk between SigB and PrfA in Listeria monocytogenes Facilitates Transitions between Extra- and Intracellular Environments. Microbiol. Mol. Biol. Rev. 83, 10.1128/mmbr.00034-19 (2019).

9. Hafner, L. et al. Differential stress responsiveness determines intraspecies virulence heterogeneity and host adaptation in Listeria monocytogenes. Nat. Microbiol. 1–17 (2024) doi:10.1038/s41564-024-01859-8.

10. Toledo-Arana, A. et al. The Listeria transcriptional landscape from saprophytism to virulence. Nature 459, 950–956 (2009).

11. Freitag, N. E., Rong, L. & Portnoy, D. A. Regulation of the prfA transcriptional activator of Listeria monocytogenes: multiple promoter elements contribute to intracellular growth and cell-to-cell spread. Infect. Immun. 61, 2537–2544 (1993).

12. Mengaud, J. et al. Pleiotropic control of Listeria monocytogenes virulence factors by a gene that is autoregulated. Mol. Microbiol. 5, 2273–2283 (1991).

13. Engelbrecht, F. et al. A new PrfA-regulated gene of encoding a small, secreted protein which belongs to the family of internalins. Mol. Microbiol. 21, 823–837 (1996).

14. Gedde, M. M., Higgins, D. E., Tilney, L. G. & Portnoy, D. A. Role of Listeriolysin O in Cell-to-Cell Spread of Listeria monocytogenes. Infect. Immun. 68, 999–1003 (2000).

15. Pistor, S., Chakraborty, T., Niebuhr, K., Domann, E. & Wehland, J. The ActA protein of Listeria monocytogenes acts as a nucleator inducing reorganization of the actin cytoskeleton. EMBO J. 13, 758–763 (1994).

16. Smith, G. A. et al. The two distinct phospholipases C of Listeria monocytogenes have overlapping roles in escape from a vacuole and cell-to-cell spread. Infect. Immun. 63, 4231–4237 (1995).

17. Vázquez-Boland, J. A. et al. Listeria Pathogenesis and Molecular Virulence Determinants. Clin. Microbiol. Rev. 14, 584 (2001).

18. Antal, E., Løberg, E., Dietrichs, E. & Mæhlen, J. Neuropathological Findings in 9 Cases of Listeria Monocytogenes Brain Stem Encephalitis. Brain Pathol. 15, 187–191 (2005).

19. Charlier, C. et al. Clinical features and prognostic factors of listeriosis: the MONALISA national prospective cohort study. Lancet Infect. Dis. 17, 510–519 (2017).

20. Oevermann, A. et al. Neuropathogenesis of Naturally Occurring Encephalitis Caused by Listeria monocytogenes in Ruminants. 10.1111/j.1750-3639.2009.00292.x (2009) doi:10.1111/j.1750-3639.2009.00292.x.

21. Henke, D. et al. Listeria monocytogenes Spreads within the Brain by Actin-Based Intra-Axonal Migration. 10.1128/IAI.00316-15 (2015) doi:10.1128/IAI.00316-15.

22. Abellanas, M. A., Purnapatre, M., Burgaletto, C. & Schwartz, M. Monocyte-derived macrophages act as reinforcements when microglia fall short in Alzheimer’s disease. Nat. Neurosci. 1–10 (2025) doi:10.1038/s41593-024-01847-5.

23. Mundt, S., Greter, M. & Becher, B. The CNS mononuclear phagocyte system in health and disease. Neuron 110, 3497–3512 (2022).

24. Frande-Cabanes, E. et al. Dissociation of Innate Immune Responses in Microglia Infected with Listeria monocytogenes. Glia 62, 233 (2014).

25. Maudet, C. et al. Bacterial inhibition of Fas-mediated killing promotes neuroinvasion and persistence. Nature 603, 900–906 (2022).

26. Birmingham, C. L. et al. Listeria monocytogenes evades killing by autophagy during colonization of host cells. Autophagy 3, 442–451 (2007).

27. Jones, G. S. & D’Orazio, S. E. F. Monocytes Are the Predominant Cell Type Associated with Listeria monocytogenes in the Gut, but They Do Not Serve as an Intracellular Growth Niche. J. Immunol. Baltim. Md 1950 198, 2796–2804 (2017).

28. Drevets, D. A. et al. The Ly-6Chigh monocyte subpopulation transports Listeria monocytogenes into the brain during systemic infection of mice. J. Immunol. Baltim. Md 1950 172, 4418–4424 (2004).

29. Bagatella, S. et al. Bovine neutrophil chemotaxis to Listeria monocytogenes in neurolisteriosis depends on microglia-released rather than bacterial factors. J. Neuroinflammation 19, (2022).

30. Dramsi, S., Lévi, S., Triller, A. & Cossart, P. Entry of Listeria monocytogenes into Neurons Occurs by Cell-to-Cell Spread: an In Vitro Study. Infect. Immun. 66, 4461–4468 (1998).

31. Rupp, S. et al. A naturally occurring prfA truncation in a Listeria monocytogenes field strain contributes to reduced replication and cell-to-cell spread. Vet. Microbiol. 179, 91–101 (2015).

32. Guldimann, C. et al. Increased spread and replication efficiency of Listeria monocytogenes in organotypic brain-slices is related to multilocus variable number of tandem repeat analysis (MLVA) complex Microbe-host interactions and microbial pathogenicity. BMC Microbiol. 15, (2015).

33. Tavares-Gomes, L. et al. Divergent host-pathogen interactions in neurolisteriosis: cytosolic replication vs. phagosomal dormancy of Listeria monocytogenes in CNS macrophages. Acta Neuropathol. (Berl.) 149, 63 (2025).

34. Kortebi, M. et al. Listeria monocytogenes switches from dissemination to persistence by adopting a vacuolar lifestyle in epithelial cells. PLOS Pathog. 13, e1006734 (2017).

35. Lotoux, A., Milohanic, E. & Bierne, H. The Viable But Non-Culturable State of Listeria monocytogenes in the One-Health Continuum. Front. Cell. Infect. Microbiol. 12, 849915 (2022).

36. Bagatella, S. et al. Intravacuolar persistence in neutrophils facilitates Listeria monocytogenes spread to co-cultured cells. mBio 0, e02700–24 (2025).

37. Helaine, S. et al. Internalization of Salmonella by Macrophages Induces Formation of Nonreplicating Persisters. Science 343, 204–208 (2014).

38. Liu, B. et al. Direct ferrous sulfate exposure facilitates the VBNC state formation rather than ferroptosis in *Listeria monocytogenes*. Microbiol. Res. 269, 127304 (2023).

39. Westermann, A. J., Barquist, L. & Vogel, J. Resolving host–pathogen interactions by dual RNA-seq. PLOS Pathog. 13, e1006033 (2017).

40. Westermann, A. J. & Vogel, J. Cross-species RNA-seq for deciphering host–microbe interactions. Nat. Rev. Genet. 22, 361–378 (2021).

41. Pisu, D., Huang, L., Grenier, J. K. & Russell, D. G. Dual RNA-Seq of Mtb-Infected Macrophages In Vivo Reveals Ontologically Distinct Host-Pathogen Interactions. Cell Rep. 30, 335–350.e4 (2020).

42. Humphrys, M. S. et al. Simultaneous Transcriptional Profiling of Bacteria and Their Host Cells. PLoS ONE 8, e80597 (2013).

43. Rienksma, R. A. et al. Comprehensive insights into transcriptional adaptation of intracellular mycobacteria by microbe-enriched dual RNA sequencing. BMC Genomics 16, 34 (2015).

44. Baddal, B. et al. Dual RNA-seq of Nontypeable Haemophilus influenzae and Host Cell Transcriptomes Reveals Novel Insights into Host-Pathogen Cross Talk. mBio 6, e01765–01715 (2015).

45. Avraham, R. et al. A highly multiplexed and sensitive RNA-seq protocol for simultaneous analysis of host and pathogen transcriptomes. Nat. Protoc. 11, 1477–1491 (2016).

46. Westermann, A. J. et al. Dual RNA-seq unveils noncoding RNA functions in host–pathogen interactions. Nature 529, 496–501 (2016).

47. Minhas, V. et al. In vivo dual RNA-seq reveals that neutrophil recruitment underlies differential tissue tropism of Streptococcus pneumoniae. *Commun*. Biol. 3, 1–12 (2020).

48. Aprianto, R., Slager, J., Holsappel, S. & Veening, J.-W. Time-resolved dual RNA-seq reveals extensive rewiring of lung epithelial and pneumococcal transcriptomes during early infection. Genome Biol. 17, 198 (2016).

49. Chatterjee, S. S. et al. Intracellular Gene Expression Profile of Listeria monocytogenes. Infect. Immun. 74, 1323 (2006).

50. Spiteri, A. G., Wishart, C. L., Pamphlett, R., Locatelli, G. & King, N. J. C. Microglia and monocytes in inflammatory CNS disease: integrating phenotype and function. Acta Neuropathol. (Berl.) 143, 179–224 (2022).

51. Myers, J. T., Tsang, A. W. & Swanson, J. A. Localized reactive oxygen and nitrogen intermediates inhibit escape of Listeria monocytogenes from vacuoles in activated macrophages. J. Immunol. Baltim. Md 1950 171, 5447–5453 (2003).

52. Schultze, T. et al. A detailed view of the intracellular transcriptome of Listeria monocytogenes in murine macrophages using RNA-seq. Front. Microbiol. 6, 1199 (2015).

53. Oliveira, A. H. et al. The Virulence and Infectivity of Listeria monocytogenes Are Not Substantially Altered by Elevated SigB Activity. Infect. Immun. 91, e00571–22.

54. Freeman, M. J., Eral, N. J. & Sauer, J.-D. Listeria monocytogenes requires phosphotransferase systems to facilitate intracellular growth and virulence. PLOS Pathog. 21, e1012492 (2025).

55. Freitag, N. E., Port, G. C. & Miner, M. D. Listeria monocytogenes — from saprophyte to intracellular pathogen. Nat. Rev. Microbiol. 7, 623–628 (2009).

56. Reniere, M. L. et al. Glutathione activates virulence gene expression of an intracellular pathogen. Nature 517, 170–173 (2015).

57. Haber, A. et al. L-glutamine Induces Expression of Listeria monocytogenes Virulence Genes. PLoS Pathog. 13, e1006161 (2017).

58. Joseph, B. et al. Identification of Listeria monocytogenes genes contributing to intracellular replication by expression profiling and mutant screening. J. Bacteriol. 188, 556–568 (2006).

59. Milohanic, E. et al. Transcriptome analysis of Listeria monocytogenes identifies three groups of genes differently regulated by PrfA. Mol. Microbiol. 47, 1613–1625 (2003).

60. Chico-Calero, I. et al. Hpt, a bacterial homolog of the microsomal glucose- 6-phosphate translocase, mediates rapid intracellular proliferation in Listeria. Proc. Natl. Acad. Sci. U. S. A. 99, 431–436 (2002).

61. Friedman, S. et al. Metabolic Genetic Screens Reveal Multidimensional Regulation of Virulence Gene Expression in Listeria monocytogenes and an Aminopeptidase That Is Critical for PrfA Protein Activation. Infect. Immun. 85, e00027–17 (2017).

62. Kunst, F. et al. The complete genome sequence of the gram-positive bacterium Bacillus subtilis. Nature 390, 249–256 (1997).

63. Narayanan, L. et al. Identification of genetic elements required for Listeria monocytogenes growth under limited nutrient conditions and virulence by a screening of transposon insertion library. Front. Microbiol. 13, 1007657 (2022).

64. Turnbough, C. L. & Switzer, R. L. Regulation of Pyrimidine Biosynthetic Gene Expression in Bacteria: Repression without Repressors. Microbiol. Mol. Biol. Rev. MMBR 72, 266–300 (2008).

65. Flåtten, I. et al. Phenotypes of dnaXE145A Mutant Cells Indicate that the Escherichia coli Clamp Loader Has a Role in the Restart of Stalled Replication Forks. J. Bacteriol. 199, e00412–17 (2017).

66. Tashjian, T. F. & Chien, P. Clamp Loader Processing Is Important during DNA Replication Stress. J. Bacteriol. 205, e00437–22.

67. Aké, F. M. D., Joyet, P., Deutscher, J. & Milohanic, E. Mutational analysis of glucose transport regulation and glucose-mediated virulence gene repression in Listeria monocytogenes. Mol. Microbiol. 81, 274–293 (2011).

68. Chen, G. Y., Pensinger, D. A. & Sauer, J.-D. Listeria monocytogenes cytosolic metabolism promotes replication, survival, and evasion of innate immunity. Cell. Microbiol. 19, e12762 (2017).

69. Stoll, R. & Goebel, W. The major PEP-phosphotransferase systems (PTSs) for glucose, mannose and cellobiose of Listeria monocytogenes, and their significance for extra- and intracellular growth. Microbiol. Read. Engl. 156, 1069–1083 (2010).

70. Tezuka, T. & Ohnishi, Y. Two Glycine Riboswitches Activate the Glycine Cleavage System Essential for Glycine Detoxification in Streptomyces griseus. J. Bacteriol. 196, 1369–1376 (2014).

71. Thomason, L. C., Court, D. L., Datta, A. R., Khanna, R. & Rosner, J. L. Identification of the Escherichia coli K-12 ybhE Gene as pgl, Encoding 6-Phosphogluconolactonase. J. Bacteriol. 186, 8248–8253 (2004).

72. Kim, K.-H. et al. Crystal structures of Enoyl-ACP reductases I (FabI) and III (FabL) from B. subtilis. J. Mol. Biol. 406, 403–415 (2011).

73. Casey, A. et al. Transcriptome analysis of Listeria monocytogenes exposed to biocide stress reveals a multi-system response involving cell wall synthesis, sugar uptake, and motility. Front. Microbiol. 5, 68 (2014).

74. Mellin, J. R. et al. Riboswitches. Sequestration of a two-component response regulator by a riboswitch-regulated noncoding RNA. Science 345, 940–943 (2014).

75. Mraheil, M. A. et al. The intracellular sRNA transcriptome of Listeria monocytogenes during growth in macrophages. Nucleic Acids Res. 39, 4235–4248 (2011).

76. Waters, L. S. & Storz, G. Regulatory RNAs in bacteria. Cell 136, 615–628 (2009).

77. Green, N. J., Grundy, F. J. & Henkin, T. M. The T box mechanism: tRNA as a regulatory molecule. FEBS Lett. 584, 318–324 (2010).

78. Zhang, J. & Ferré-D’Amaré, A. R. Structure and mechanism of the T-box riboswitches. Wiley Interdiscip. Rev. RNA 6, 419–433 (2015).

79. Izar, B., Mraheil, M. A. & Hain, T. Identification and Role of Regulatory Non-Coding RNAs in Listeria monocytogenes. Int. J. Mol. Sci. 12, 5070–5079 (2011).

80. Dar, D. et al. Term-seq reveals abundant ribo-regulation of antibiotics resistance in bacteria. Science 352, aad9822 (2016).

81. Duru, I. C. et al. High-pressure processing-induced transcriptome response during recovery of Listeria monocytogenes. BMC Genomics 22, 117 (2021).

82. Camejo, A. et al. In vivo transcriptional profiling of Listeria monocytogenes and mutagenesis identify new virulence factors involved in infection. PLoS Pathog. 5, e1000449 (2009).

83. Morawska, L. P. & Kuipers, O. P. Transcriptome analysis and prediction of the metabolic state of stress-induced viable but non-culturable Bacillus subtilis cells. Sci. Rep. 12, 18015 (2022).

84. Tavares-Gomes, L. et al. Transcriptome of microglia reveals a species-specific expression profile in bovines with conserved and new signature genes. Glia 69, 1932–1949 (2021).

85. Hsieh, C. S. et al. Development of TH1 CD4+ T cells through IL-12 produced by Listeria-induced macrophages. Science 260, 547–549 (1993).

86. McKenna, S. et al. Endotoxemia Induces IκBβ/NF-κB-Dependent Endothelin-1 Expression in Hepatic Macrophages. J. Immunol. Baltim. Md 1950 195, 3866–3879 (2015).

87. Romani, L., Puccetti, P. & Bistoni, F. Interleukin-12 in infectious diseases. Clin. Microbiol. Rev. 10, 611–636 (1997).

88. Wagner, R. D., Steinberg, H., Brown, J. F. & Czuprvnski, C. J. Recombinant interleukin-12 enhances resistance of mice to *Listeria monocytogenes* infection. Microb. Pathog. 17, 175–186 (1994).

89. Wahl, J. R. et al. Murine Macrophages Produce Endothelin-1 After Microbial Stimulation. Exp. Biol. Med. 230, 652–658 (2005).

90. Ding, G. et al. Porcine Reproductive and Respiratory Syndrome Virus Structural Protein GP3 Regulates Claudin 4 To Facilitate the Early Stages of Infection. J. Virol. 94, e00124–20 (2020).

91. Ji, W., Zhuang, X., Jiang, W. G. & Martin, T. A. Tight junctional protein family, Claudins in cancer and cancer metastasis. Front. Oncol. 15, (2025).

92. Ganesan, S. & Roy, C. R. Host cell depletion of tryptophan by IFNγ-induced Indoleamine 2,3-dioxygenase 1 (IDO1) inhibits lysosomal replication of Coxiella burnetii. PLOS Pathog. 15, e1007955 (2019).

93. Niño-Castro, A. et al. The IDO1-induced kynurenines play a major role in the antimicrobial effect of human myeloid cells against Listeria monocytogenes. Innate Immun. 20, 401–411 (2014).

94. Popov, A. et al. Indoleamine 2,3-dioxygenase–expressing dendritic cells form suppurative granulomas following *Listeria monocytogenes* infection. J. Clin. Invest. 116, 3160–3170 (2006).

95. Brunetta, E. et al. Macrophage expression and prognostic significance of the long pentraxin PTX3 in COVID-19. Nat. Immunol. 22, 19–24 (2021).

96. Jeon, H., Lee, S., Lee, W.-H. & Suk, K. Analysis of glial secretome: The long pentraxin PTX3 modulates phagocytic activity of microglia. J. Neuroimmunol. 229, 63–72 (2010).

97. Villar, J., Ouaknin, L., Cros, A. & Segura, E. Monocytes differentiate along two alternative pathways during sterile inflammation. EMBO Rep. 24, e56308 (2023).

98. Singh, R., Jamieson, A. & Cresswell, P. GILT is a critical host factor for Listeria monocytogenes infection. Nature 455, 1244–1247 (2008).

99. Satoh, J.-I., Kino, Y., Yanaizu, M., Ishida, T. & Saito, Y. Microglia express gamma-interferon-inducible lysosomal thiol reductase in the brains of Alzheimer’s disease and Nasu-Hakola disease. Intractable Rare Dis. Res. 7, 251–257 (2018).

100. Galli, G. & Saleh, M. Immunometabolism of Macrophages in Bacterial Infections. Front. Cell. Infect. Microbiol. 10, (2021).

101. Rosenberg, G., Riquelme, S., Prince, A. & Avraham, R. Immunometabolic crosstalk during bacterial infection. Nat. Microbiol. 7, 497–507 (2022).

102. Cabal-Hierro, L. et al. TRAF-mediated modulation of NF-kB AND JNK activation by TNFR2. Cell. Signal. 26, 2658–2666 (2014).

103. Devergne, O. et al. Association of TRAF1, TRAF2, and TRAF3 with an Epstein-Barr virus LMP1 domain important for B-lymphocyte transformation: role in NF-kappaB activation. Mol. Cell. Biol. 16, 7098–7108 (1996).

104. Xie, P. TRAF molecules in cell signaling and in human diseases. J. Mol. Signal. 8, 7 (2013).

105. Gupta, S. et al. IL-6 augments IL-4-induced polarization of primary human macrophages through synergy of STAT3, STAT6 and BATF transcription factors. Oncoimmunology 7, e1494110 (2018).

106. Sahoo, A. et al. Batf is important for IL-4 expression in T follicular helper cells. Nat. Commun. 6, 7997 (2015).

107. Wculek, S. K., Dunphy, G., Heras-Murillo, I., Mastrangelo, A. & Sancho, D. Metabolism of tissue macrophages in homeostasis and pathology. Cell. Mol. Immunol. 19, 384–408 (2022).

108. Bathke, J., Konzer, A., Remes, B., McIntosh, M. & Klug, G. Comparative analyses of the variation of the transcriptome and proteome of Rhodobacter sphaeroides throughout growth. BMC Genomics 20, 358 (2019).

109. Ghazalpour, A. et al. Comparative Analysis of Proteome and Transcriptome Variation in Mouse. PLoS Genet. 7, e1001393 (2011).

110. Boddaert, J. et al. CD8 signaling in microglia/macrophage M1 polarization in a rat model of cerebral ischemia. PloS One 13, e0186937 (2018).

111. Bruce, K. D. Lipoprotein Lipase Regulates Lipid and Lipoprotein Processing in Microglia. Alzheimers Dement. 18, e060210 (2022).

112. Loving, B. A. et al. Lipoprotein Lipase Regulates Microglial Lipid Droplet Accumulation. Cells 10, 198 (2021).

113. Mangale, V. et al. Microglia influence host defense, disease, and repair following murine coronavirus infection of the central nervous system. Glia 68, 2345–2360 (2020).

114. Sousa, C. et al. Single-cell transcriptomics reveals distinct inflammation-induced microglia signatures. EMBO Rep. 19, e46171 (2018).

115. Arnett, E. et al. The Pore-Forming Toxin Listeriolysin O Is Degraded by Neutrophil Metalloproteinase-8 and Fails To Mediate Listeria monocytogenes Intracellular Survival in Neutrophils. J. Immunol. Author Choice 192, 234–244 (2014).

116. Gluschko, A. et al. Macrophages target Listeria monocytogenes by two discrete non-canonical autophagy pathways. Autophagy 18, 1090–1107 (2022).

117. Palma, S. D. et al. Comparative spatiotemporal analysis of the intrathecal immune response in natural listeric rhombencephalitis of cattle and small ruminants. Comp. Immunol. Microbiol. Infect. Dis. 35, 429–441 (2012).

118. Burke, T. P. et al. Listeria monocytogenes Is Resistant to Lysozyme through the Regulation, Not the Acquisition, of Cell Wall-Modifying Enzymes. J. Bacteriol. 196, 3756–3767 (2014).

119. Rae, C. S., Geissler, A., Adamson, P. C. & Portnoy, D. A. Mutations of the Listeria monocytogenes Peptidoglycan N-Deacetylase and O-Acetylase Result in Enhanced Lysozyme Sensitivity, Bacteriolysis, and Hyperinduction of Innate Immune Pathways ▿. Infect. Immun. 79, 3596–3606 (2011).

120. Aguilar-Bultet, L. et al. Genetic separation of Listeria monocytogenes causing central nervous system infections in animals. Front. Cell. Infect. Microbiol. 8, (2018).

121. Dreyer, M. et al. Listeria monocytogenes sequence type 1 is predominant in ruminant rhombencephalitis. Sci. Rep. 6, 36419 (2016).

122. Blériot, C. et al. Liver-Resident Macrophage Necroptosis Orchestrates Type 1 Microbicidal Inflammation and Type-2-Mediated Tissue Repair during Bacterial Infection. Immunity 42, 145–158 (2015).

123. Nowacki, J. S., Jones, G. S. & D’Orazio, S. E. F. Listeria monocytogenes use multiple mechanisms to disseminate from the intestinal lamina propria to the mesenteric lymph nodes. Microbiol. Spectr. 0, e02595–24 (2024).

124. Bogdan, C. Macrophages as host, effector and immunoregulatory cells in leishmaniasis: Impact of tissue micro-environment and metabolism. Cytokine X 2, 100041 (2020).

125. Hoffman, D. et al. A non-classical monocyte-derived macrophage subset provides a splenic replication niche for intracellular *Salmonella*. Immunity 54, 2712–2723.e6 (2021).

126. Leiba, J. et al. Dynamics of macrophage polarization support Salmonella persistence in a whole living organism. eLife 13, e89828 (2024).

127. Pessenda, G. et al. Kupffer cell and recruited macrophage heterogeneity orchestrate granuloma maturation and hepatic immunity in visceral leishmaniasis. Nat. Commun. 16, 3125 (2025).

128. Fischer, M. A. et al. Listeria monocytogenes genes supporting growth under standard laboratory cultivation conditions and during macrophage infection. Genome Res. 32, 1711–1726 (2022).

129. Chambers, K. A. & Scheck, R. A. Bacterial virulence mediated by orthogonal post-translational modification. Nat. Chem. Biol. 16, 1043–1051 (2020).

130. Hommel, B. et al. Cryptococcus neoformans resists to drastic conditions by switching to viable but non-culturable cell phenotype. PLOS Pathog. 15, e1007945 (2019).

131. . Lefrançois, L. H., et al. Temporal genome-wide fitness analysis of Mycobacterium marinum during infection reveals the genetic requirement for virulence and survival in amoebae and microglial cells. mSystems 9, e0132623 (2024).

132. Rytter, H. et al. Dual proteomics of infected macrophages reveal bacterial and host players involved in the Francisella intracellular life cycle and cell to cell dissemination by merocytophagy. Sci. Rep. 14, 7797 (2024).

133. Telser, J. et al. Metabolic reprogramming of Salmonella infected macrophages and its modulation by iron availability and the mTOR pathway. Microb. Cell 6, 531–543.

134. Wagner, E. G. H. & Romby, P. Small RNAs in bacteria and archaea: who they are, what they do, and how they do it. Adv. Genet. 90, 133–208 (2015).

135. Molmeret, M., Horn, M., Wagner, M., Santic, M. & Abu Kwaik, Y. Amoebae as Training Grounds for Intracellular Bacterial Pathogens. Appl. Environ. Microbiol. 71, 20–28 (2005).

136. Daniel, J., Maamar, H., Deb, C., Sirakova, T. D. & Kolattukudy, P. E. Mycobacterium tuberculosis uses host triacylglycerol to accumulate lipid droplets and acquires a dormancy-like phenotype in lipid-loaded macrophages. PLoS Pathog. 7, e1002093 (2011).

137. Daniel, J. et al. Induction of a novel class of diacylglycerol acyltransferases and triacylglycerol accumulation in Mycobacterium tuberculosis as it goes into a dormancy-like state in culture. J. Bacteriol. 186, 5017–5030 (2004).

138. Galagan, J. E. et al. The Mycobacterium tuberculosis regulatory network and hypoxia. Nature 499, 178–183 (2013).

139. Carvalho, F. et al. Aquatic environment drives the emergence of cell wall-deficient dormant forms in Listeria. Nat. Commun. 15, 8499 (2024).

140. Liu, B. et al. Direct ferrous sulfate exposure facilitates the VBNC state formation rather than ferroptosis in *Listeria monocytogenes*. Microbiol. Res. 269, 127304 (2023).

141. Crimmins, G. T. et al. Listeria monocytogenes multidrug resistance transporters activate a cytosolic surveillance pathway of innate immunity. Proc. Natl. Acad. Sci. 105, 10191–10196 (2008).

142. Kaplan Zeevi, M., et al. Listeria monocytogenes Multidrug Resistance Transporters and Cyclic Di-AMP, Which Contribute to Type I Interferon Induction, Play a Role in Cell Wall Stress. J. Bacteriol. 195, 5250–5261 (2013).

143. Vimberg, V., Zieglerová, L., Buriánková, K., Branny, P. & Balíková Novotná, G. VanZ Reduces the Binding of Lipoglycopeptide Antibiotics to Staphylococcus aureus and Streptococcus pneumoniae Cells. Front. Microbiol. 11, 566 (2020).

144. Wielders, C. L. C., Fluit, A. C., Brisse, S., Verhoef, J. & Schmitz, F. J. mecA Gene Is Widely Disseminated in Staphylococcus aureus Population. J. Clin. Microbiol. 40, 3970–3975 (2002).

145. Tanaka, N. & Shuman, S. RtcB Is the RNA Ligase Component of an Escherichia coli RNA Repair Operon. J. Biol. Chem. 286, 7727–7731 (2011).

146. König, P. et al. The VBNC state: a fundamental survival strategy of Acinetobacter baumannii. mBio 14, e02139–23.

147. Peyrusson, F. et al. Intracellular Staphylococcus aureus persisters upon antibiotic exposure. Nat. Commun. 11, 2200 (2020).

148. Amici, S. A. et al. CD38 Is Robustly Induced in Human Macrophages and Monocytes in Inflammatory Conditions. Front. Immunol. 9, 1593 (2018).

149. Lischke, T. et al. CD38 controls the innate immune response against Listeria monocytogenes. Infect. Immun. 81, 4091–4099 (2013).

150. Osborne, S. E. et al. Type I interferon promotes cell-to-cell spread of Listeria monocytogenes. Cell. Microbiol. 19, (2017).

151. Dussurget, O., Bierne, H. & Cossart, P. The bacterial pathogen Listeria monocytogenes and the interferon family: type I, type II and type III interferons. Front. Cell. Infect. Microbiol. 4, (2014).

152. Italiani, P. & Boraschi, D. From Monocytes to M1/M2 Macrophages: Phenotypical vs. Functional Differentiation. Front. Immunol. 5, 514 (2014).

153. Lannes, N. et al. Interactions of human microglia cells with Japanese encephalitis virus. Virol. J. 14, 8 (2017).

154. Boneca, I. G. et al. A critical role for peptidoglycan N-deacetylation in Listeria evasion from the host innate immune system. Proc. Natl. Acad. Sci. 104, 997–1002 (2007).

155. Monk, I. R., Gahan, C. G. M. & Hill, C. Tools for Functional Postgenomic Analysis of Listeria monocytogenes. Appl. Environ. Microbiol. 74, 3921–3934 (2008).

156. Cingolani, P. et al. A program for annotating and predicting the effects of single nucleotide polymorphisms, SnpEff: SNPs in the genome of Drosophila melanogaster strain w1118; iso-2; iso-3. Fly (Austin) 6, 80–92 (2012).

157. Gözel, B. et al. Hyperinvasiveness of Listeria monocytogenes sequence type 1 is independent of lineage I-specific genes encoding internalin-like proteins. 10.1002/mbo3.790 (2019) doi:10.1002/mbo3.790.

158. Pownall, W. R. et al. Safety of a Novel Listeria monocytogenes-Based Vaccine Vector Expressing NcSAG1 (Neospora caninum Surface Antigen 1). Front. Cell. Infect. Microbiol. 11, 675219 (2021).

159. Villa-Rodríguez, E., Ibarra-Gámez, C. & de Los Santos-Villalobos, S. Extraction of high-quality RNA from Bacillus subtilis with a lysozyme pre-treatment followed by the Trizol method. J. Microbiol. Methods 147, 14–16 (2018).

160. Andrews S 2010. FastQC: a quality control tool for high throughput sequence data. https://www.bioinformatics.babraham.ac.uk/projects/fastqc/.

161. Wang, L., Wang, S. & Li, W. RSeQC: quality control of RNA-seq experiments. Bioinforma. Oxf. Engl. 28, 2184–2185 (2012).

162. Kim, D., Langmead, B. & Salzberg, S. L. HISAT: a fast spliced aligner with low memory requirements. Nat. Methods 12, 357–360 (2015).

163. Liao, Y., Smyth, G. K. & Shi, W. featureCounts: an efficient general purpose program for assigning sequence reads to genomic features. Bioinforma. Oxf. Engl. 30, 923–930 (2014).

164. Love, M. I., Huber, W. & Anders, S. Moderated estimation of fold change and dispersion for RNA-seq data with DESeq2. Genome Biol. 15, 550 (2014).

165. Subramanian, A. et al. Gene set enrichment analysis: a knowledge-based approach for interpreting genome-wide expression profiles. Proc. Natl. Acad. Sci. U. S. A. 102, 15545–15550 (2005).

166. Wu, T. et al. clusterProfiler 4.0: A universal enrichment tool for interpreting omics data. Innov. Camb. Mass 2, 100141 (2021).

167. Kanehisa, M., Sato, Y. & Kawashima, M. KEGG mapping tools for uncovering hidden features in biological data. Protein Sci. Publ. Protein Soc. 31, 47–53 (2022).

168. Liberzon, A. et al. The Molecular Signatures Database (MSigDB) hallmark gene set collection. Cell Syst. 1, 417–425 (2015).

169. Memedovski, R. et al. Investigation of the mechanism of action of mefloquine and derivatives against the parasite Echinococcus multilocularis. Int. J. Parasitol. Drugs Drug Resist. 21, 114–124 (2023).

170. UniProt Consortium. UniProt: a worldwide hub of protein knowledge. Nucleic Acids Res. 47, D506–D515 (2019).

171. Semeraro, M. et al. Transient Adaptation of Toxoplasma gondii to Exposure by Thiosemicarbazone Drugs That Target Ribosomal Proteins Is Associated with the Upregulated Expression of Tachyzoite Transmembrane Proteins and Transporters. Int. J. Mol. Sci. 25, 9067 (2024).

172. Yu, F. et al. Fast Quantitative Analysis of timsTOF PASEF Data with MSFragger and IonQuant. Mol. Cell. Proteomics MCP 19, 1575–1585 (2020).

173. Haider, S. & Pal, R. Integrated Analysis of Transcriptomic and Proteomic Data. Curr. Genomics 14, 91–110 (2013).

